# Membrane Lipids Augment Cell Envelope Stress Signaling and Resistance to Antibiotics and Antimicrobial Peptides in *Enterococcus faecalis*

**DOI:** 10.1101/2023.10.17.562839

**Authors:** William R Miller, April Nguyen, Kavindra V Singh, Samie Rizvi, Ayesha Khan, Sam G Erickson, Stephanie L Egge, Melissa Cruz, An Q Dinh, Lorena Diaz, Rutan Zhang, Libin Xu, Danielle A Garsin, Yousif Shamoo, Cesar A Arias

**Affiliations:** Division of Infectious Diseases, Houston Methodist Hospital, Houston, Texas, USA; Center for Infectious Diseases, Houston Methodist Research Institute, Houston, Texas, USA; Department of Medicine, Weill Cornell Medical College, New York, New York, USA; McGovern Medical School, University of Texas Health Science Center, Houston, TX, USA; Microbiology and Molecular Genetics, Graduate School of Biomedical Sciences, University of Texas Health Science Center, Houston, TX, USA; Molecular Genetics and Antimicrobial Resistance Unit, Universidad El Bosque, Bogota, Colombia; Genomics and Resistant Microbes Group, Facultad de Medicina Clinica Alemana, Universidad del Desarrollo and Millennium Initiative for Collaborative Research On Bacterial Resistance (MICROB-R), Santiago, Chile; Department of Medicinal Chemistry, University of Washington, Seattle, Washington, USA; Department of Biosciences, Rice University, Houston, TX, USA

## Abstract

Enterococci have evolved resistance mechanisms to protect their cell envelopes against bacteriocins and host cationic antimicrobial peptides (CAMPs) produced in the gastrointestinal environment. Activation of the membrane stress response has also been tied to resistance to the lipopeptide antibiotic daptomycin. However, the actual effectors mediating resistance have not been elucidated. Here, we show that the MadRS (formerly YxdJK) membrane antimicrobial peptide defense system controls a network of genes, including a previously uncharacterized three gene operon (*madEFG*) that protects the *E. faecalis* cell envelope from antimicrobial peptides. Constitutive activation of the system confers protection against CAMPs and daptomycin in the absence of a functional LiaFSR system and leads to persistence of cardiac microlesions *in vivo*. Moreover, changes in the lipid cell membrane environment alter CAMP susceptibility and expression of the MadRS system. Thus, we provide a framework supporting a multilayered envelope defense mechanism for resistance and survival coupled to virulence.

---

Enterococci are a leading cause of healthcare-associated infections and possess a diverse array of intrinsic and acquired antimicrobial resistance determinants.^1,2^ These organisms are commensal colonizers of the gastrointestinal tract, where they are exposed to cationic antimicrobial peptides (CAMPs) produced by the innate immune system.^3^ Several important antibiotics used in clinical practice, including the lipopeptide daptomycin, share functional similarities with CAMPs.^4^ Daptomycin possesses a peptide core with an acyl tail, and in conjunction with calcium forms an amphipathic complex that binds the membrane lipid phosphatidylglycerol (PG) and the undecaprenyl intermediates of cell wall synthesis.^5^ Daptomycin binding displaces enzymes necessary for cell envelope genesis, ultimately leading to membrane disruption and bacterial cell death. The widespread frequency of multidrug resistance in both *Enterococcus faecium* and *E. faecalis* has led to a reliance on daptomycin for treatment of recalcitrant invasive vancomycin-resistant enterococcal (VRE) infections.

As members of the gastrointestinal flora, enterococci have evolved several mechanisms to sense and respond to CAMPs. This survival strategy involves activation of cell stress response signaling systems in conjunction with changes in the synthesis of cell membrane phospholipids.^6–9^ In particular, the LiaFSR system of *E. faecalis* has been linked to survival in the presence of daptomycin, the cathelicidin LL-37, and other CAMPs, via a secreted protein (LiaX), which plays a dual sensing and regulatory role mediated by the N-and C-terminal domains, respectively.^10^ While substitutions in the LiaFSR system lead to antibiotic tolerance, or loss of bactericidal daptomycin activity, additional changes in enzymes related to membrane lipid biosynthesis are required for the full resistance phenotype. Since the LiaFSR pathway is also a major mediator of daptomycin resistance in clinical settings, the selective pressure exerted by CAMPs has important therapeutic repercussions.

Experimental evolution in the presence of daptomycin using both *E. faecalis* and *E. faecium* strains lacking the LiaR response regulator and, therefore, deficient in LiaFSR signaling, suggested that activation of the YxdJK two-component system in the presence of mutations in *dak* (encoding a putative fatty acid kinase) could also lead to the development of daptomycin resistance.^7,11^ The YxdJK system shares homology with other ABC transporters involved in lantibiotic and antimicrobial peptide resistance, including the BceRS-BceAB system from *Bacillus subtilis* and the GraRS-VraDE system from *Staphylococcus aureus*, although it falls in a distinct group apart from these two systems based on phylogenetic analysis.^12^ Dak is a homologue of the fatty acid kinase Fak in *S. aureus*, responsible for the first step in incorporation of exogenous fatty acids into bacterial phospholipids.^13^ Of note, fatty acids have been implicated in increasing survival in the presence of daptomycin and other membrane stressors.^14^ As the specific mechanism by which *E. faecalis* defends the cell envelope against CAMPs is unknown, we sought to determine the role of *dak* and the YxdJK system (which we hereafter refer to as the MadRS system for <u>m</u>embrane <u>a</u>ntimicrobial peptide <u>d</u>efense), in cell envelope protection against daptomycin and host-derived CAMPs.

## Results

### Identification of the MadRS regulon

We previously described an alanine to aspartate substitution at amino acid position 202 in the MadS histidine kinase associated with daptomycin resistance in *E. faecalis*.^7^ To investigate the functional importance of this change, we introduced this allele into *E. faecalis* OG1RF (OG1RF*madS_A202E_*). The presence of *madS_A202E_* was sufficient to increase the minimum inhibitory concentration (MIC) of daptomycin from 1.5 to 6 μg/mL, and the MIC of bacitracin from 32 to 96 μg/mL (**Table 1**). In the OG117 background (a derivative of OG1RF adapted for CRISPR based manipulation), introduction of both the *madS_A202E_* allele and deletion of the *dak* gene, led to an increase in daptomycin and bacitracin MICs (**Table 1**).

**Table 1.**
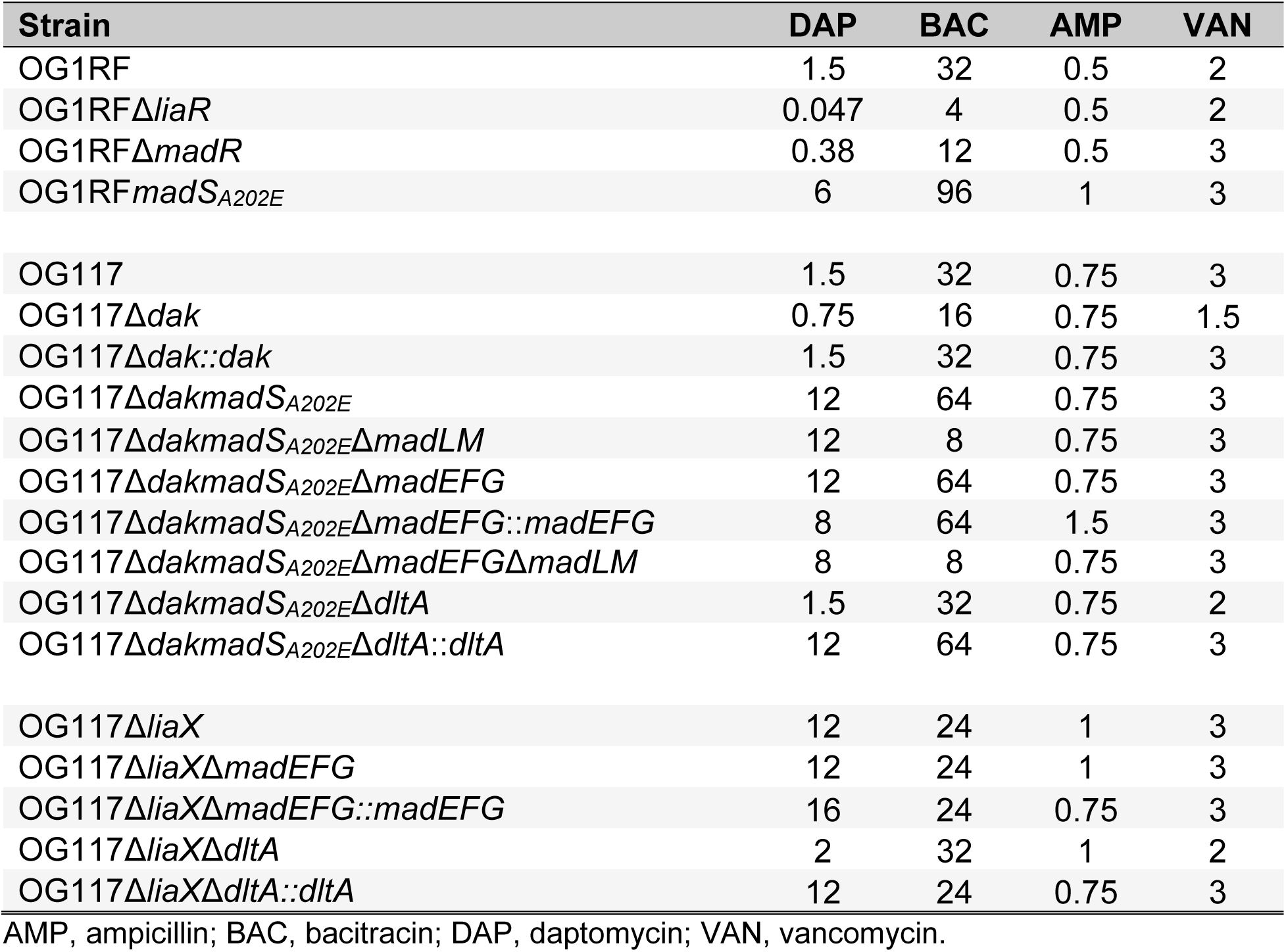
Antimicrobial Minimum Inhibitory Concentrations (all values μg/mL).

| Strain | DAP | BAC | AMP | VAN |
| --- | --- | --- | --- | --- |
| OG1RF | 1.5 | 32 | 0.5 | 2 |
| OG1RFΔ*liaR* | 0.047 | 4 | 0.5 | 2 |
| OG1RFΔ*madR* | 0.38 | 12 | 0.5 | 3 |
| OG1RF*madS*<sub>A202E</sub> | 6 | 96 | 1 | 3 |
| OG117 | 1.5 | 32 | 0.75 | 3 |
| OG117Δ*dak* | 0.75 | 16 | 0.75 | 1.5 |
| OG117Δ*dak::dak* | 1.5 | 32 | 0.75 | 3 |
| OG117Δ*dakmadS*<sub>A202E</sub> | 12 | 64 | 0.75 | 3 |
| OG117Δ*dakmadS*<sub>A202E</sub>Δ*madLM* | 12 | 8 | 0.75 | 3 |
| OG117Δ*dakmadS*<sub>A202E</sub>Δ*madEFG* | 12 | 64 | 0.75 | 3 |
| OG117Δ*dakmadS*<sub>A202E</sub>Δ*madEFG::madEFG* | 8 | 64 | 1.5 | 3 |
| OG117Δ*dakmadS*<sub>A202E</sub>Δ*madEFG*Δ*madLM* | 8 | 8 | 0.75 | 3 |
| OG117Δ*dakmadS*<sub>A202E</sub>Δ*dltA* | 1.5 | 32 | 0.75 | 2 |
| OG117Δ*dakmadS*<sub>A202E</sub>Δ*dltA::dltA* | 12 | 64 | 0.75 | 3 |
| OG117Δ*liaX* | 12 | 24 | 1 | 3 |
| OG117Δ*liaX*Δ*madEFG* | 12 | 24 | 1 | 3 |
| OG117Δ*liaX*Δ*madEFG::madEFG* | 16 | 24 | 0.75 | 3 |
| OG117Δ*liaX*Δ*dltA* | 2 | 32 | 1 | 2 |
| OG117Δ*liaX*Δ*dltA::dltA* | 12 | 24 | 0.75 | 3 |

AMP, ampicillin; BAC, bacitracin; DAP, daptomycin; VAN, vancomycin.

Although the MadRS system is known to mediate protection from the antibiotic bacitracin via the presence of the ABC transporter MadLM (previously YxdLM), we hypothesized that this transporter may also be involved in the daptomycin resistance phenotype, similar to VraDE in *S. aureus*.^15,16^ However, deletion of the genes encoding MadLM from the chromosome of the daptomycin resistant *E. faecalis* OG117Δ*dakmadS_A202E_*led to a decrease in the MIC of bacitracin, but did not alter susceptibility to daptomycin (**Table 1**). Thus, we applied a transcriptomic approach to identify additional genes under control of the MadRS system that could explain the changes in daptomycin MIC.

We compared differentially expressed genes in OG1RFΔ*madR*, which lacks the response regulator of the MadRS system, and OG1RF*madS_A202E_* to the parent strain OG1RF in the presence and absence of daptomycin (**Figure 1A-B**, **Table S1**). Comparing OG1RFΔ*madR* to OG1RF, there were 8 genes with a significant difference in expression in the absence of daptomycin exposure. The most strongly downregulated was a previously unidentified putative operon of three genes, *madEFG*, followed by *madLM*, as well as the genes *mprF2* (which catalyzes the production of lysyl-PG) and *salA* (a peptidoglycan hydrolase, **Table S1**). In OG1RF*madS_A202E_*, there were 364 and 368 differentially expressed genes in the absence and presence of daptomycin, respectively. The most significant changes included increased expression of the *madEFG* operon, the *dlt* operon (involved in D-alanylation of lipoteichoic acids), *madLM*, *salA*, and downregulation of the *upp* gene, annotated as a uracil phosphoribosyltransferase. To confirm these results, we performed qRT-PCR for the genes *madG*, *madL*, *dltA*, and *madA*, which was previously identified as a component of the MadRS signaling network with minimal differences in expression under bacitracin induction. In the *madR* knockout, a decrease in gene expression for *madG* and *madL* was identified as compared to OG1RF both in the presence and absence of daptomycin. There was no significant difference in expression of either *madA* or *dltA*. In the *madS_A202E_*background, there was a significant increase in the expression of *madG*, *madL*, and *dltA*, particularly on exposure to daptomycin (**Figure 1C**).

**Figure 1.**
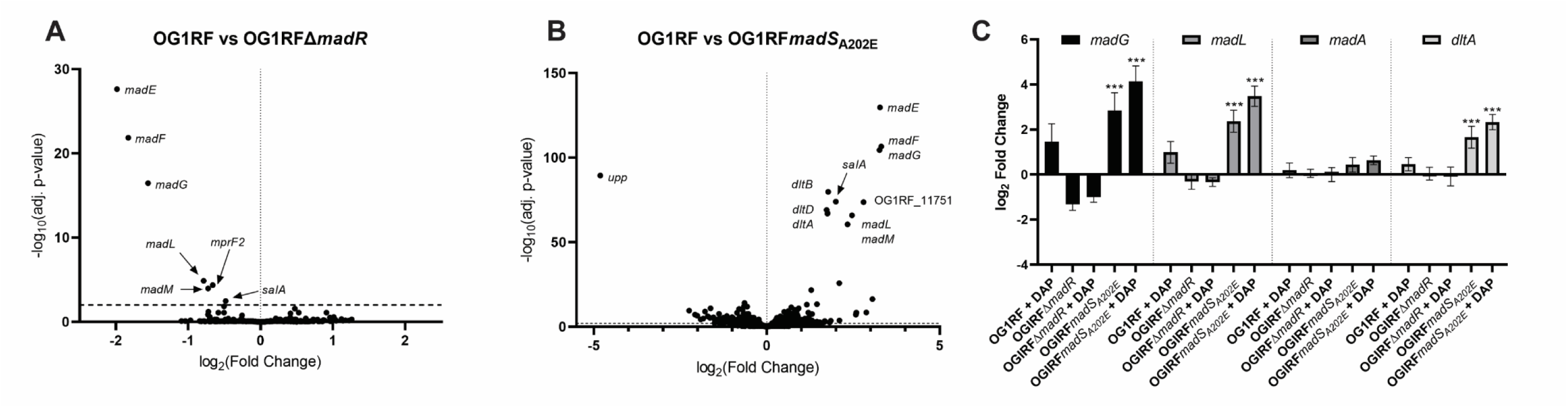
Transcriptional profile of the MadRS regulon. A) Gene expression by RNA sequence analysis of OG1RFΔ*madR* without daptomycin exposure as compared to wild type OG1RF. The log_2_ fold change in gene expression is shown on the x-axis, and -log_10_(adjusted p-value) is shown on the y-axis. Horizontal dotted line represents the significance cutoff of p < 0.01. B) Differentially expressed genes in the OG1RF*madS_A202E_* as compared to OG1RF without daptomycin exposure. Select genes are labeled on the plot, a full list of differentially expressed genes can be found in Supplemental Table 1. C) Quantitative reverse-transcriptase polymerase chain reaction qRT-PCR) of representative genes of the MadRS system. Strains and antibiotic exposure conditions are shown on the x-axis. Log_2_ fold change in gene expression is shown on the y-axis. Error bars represent the standard deviation of three independent biological runs performed in technical triplicate. Significant differences in gene expression were compared to OG1RF without exposure to daptomycin. ***, p < 0.001.

### The MadRS network provides a targeted response to antibiotics and host derived AMPs

To study the effects of each component of the MadRS system, we constructed a series of deletion mutants targeting *madLM*, *madEFG* and *dltA* in daptomycin-resistant OG117Δ*dakmadS_A202E_*. Deletion of *madLM* or *madEFG* had no impact on the MIC of daptomycin, while deletion of both led to a modest decrease (12 to 8 μg/mL; **Table 1**). In contrast, deletion of *dltA* resulted in a reduction of 8-fold in the in daptomycin MIC (12 to 1.5 μg/mL), similar to wild type OG117. Since *madEFG* did not appear to alter susceptibility to either bacitracin or daptomycin but were among the most significantly expressed genes under the control of MadR, we investigated alternative roles for MadEFG.

Given the similarities between daptomycin and CAMPs, we assayed the activity of several representative CAMPs, including LL-37 (cathelicidin), human β-defensin 3 (HBD3), and human neutrophil peptide-1 (HNP1, α-defensin) against the MadRS knockout mutants (**Figure 2A-B, Supplemental Figure 1**). In the presence of LL-37 at 50 μg/mL, there was a significant increase in survival of OG117Δ*dakmadS_A202E_* versus OG117 (mean difference 42.6%, p=0.0001). Deletion of *madEFG*, but not *madLM* or *dltA*, resulted in a reversion of the survival benefit (mean difference from OG117 1.79%, p=0.99), and complementation with *madEFG* at the native chromosomal location restored bacterial survival (mean difference 36%, p=0.0001). For HBD3, deletion of *dak* was sufficient to increase survival (mean difference 47.4%, p=0.0003), comparable to the survival of the OG117Δ*dakmadS_A202E_*mutant. Restoring the *dak* gene in the native chromosomal location restored HBD3 activity to wild type levels (mean difference 12.2%, p=0.72). Loss of *madEFG* resulted in a decrease in HBD3 survival by 10.1%, but this was not statistically significant. Interestingly, no change in survival was seen for any strain with the α-defensin HNP1 (**Supplemental Figure 1)**.

**Figure 2.**
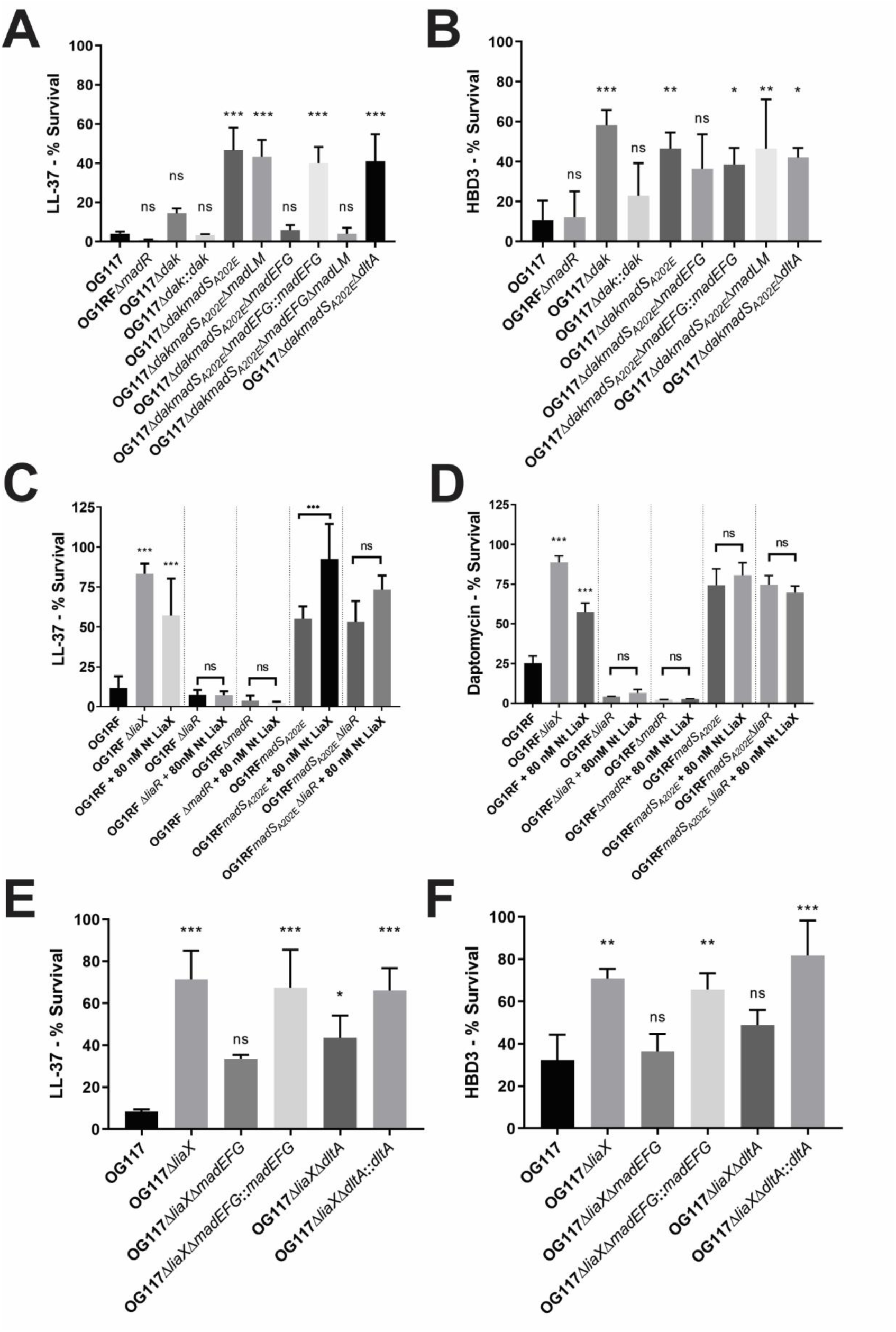
MadRS mediates survival in the presence of antimicrobial peptides and daptomycin independent of the LiaFSR system. A) LL-37 killing assay (50 µg/mL of LL-37) performed in OG117 background. Percent survival was calculated by dividing the colony forming units per milliliter (CFU/mL) of the peptide free growth control with the CFU/mL of the peptide treated samples. The OG117Δ*dakmadS_A202E_* strain (daptomycin resistant) showed a significant increase in survival as compared to wild type OG117. Deletion of *madEFG*, but not *madLM* alone or *dltA* led to increased killing by LL-37, and survival was restored on complementation of *madEFG* in the native chromosomal location. B) HBD3 killing assay (6 µg/mL of HBD3) performed in the OG117 background. Deletion of *dak* was sufficient to increase survival in the presence of HBD3. C) LL-37 and D) daptomycin (4 µg/mL) killing assay in the OG1RF background, with and without the addition of N-terminal LiaX protein (Nt LiaX), previously shown to protect *E. faecalis* from LL-37 mediated killing in a LiaFSR dependent manner. Nt LiaX was unable to rescue a MadR knockout strain (pairwise comparisons are shown as a bracket above strain pairs), and the presence of MadS_A202E_ was able to confer increased survival in the presence of both LL-37 and daptomycin in a strain lacking a functional LiaFSR system (Δ*liaR*). E) LL-37 and F) HBD3 killing assay in the OG117Δ*liaX* background with a constitutively active LiaFSR pathway. Deletion of *madEFG* decreased survival to a level that was not statistically different from wild type. In the absence of membrane changes induced by loss of *dak*, *madEFG* was also noted to be important for survival in the presence of HBD3. Significant differences in survival between each strain and OG117 was determined by one-way ANOVA. Error bars represent the standard deviation of three independent runs, performed in biological triplicate on separate days. *, p < 0.05; **, p < 0.01; ***, p < 0.001; ns, not significant.

### MadRS mediates survival in the presence of CAMPs independent of LiaFSR

The LiaFSR system has been implicated as the major mediator of cell envelope homeostasis under daptomycin stress in enterococci. Furthermore, the addition of exogenous N-terminal LiaX has been shown to activate LiaFSR signaling and provide protection against membrane stress even in daptomycin-sensitive *E. faecalis* isolates.^10^ Thus, we sought to characterize the distinct roles of MadRS compared to LiaFSR in the cell envelope stress response. Introduction of the *madS_A202E_* allele into OG1RFΔ*liaR* (lacking activation of LiaFSR due to deletion of the gene encoding the response regulator) was sufficient to increase survival in the presence of LL-37 (mean difference 41.4%, p<0.0001, **Figure 2C**), suggesting that a functional LiaFSR response was not necessary for MadRS mediated protection against LL-37. Conversely, deletion of the gene encoding the MadR response regulator in OG1RF resulted in decreased survival in the presence of LL-37 (3.8% versus 11.8%), an effect that could not be rescued with the addition of exogenous LiaX, despite the strain retaining a functional LiaFSR system. A similar pattern was seen with daptomycin, with decreased survival in the *madR* deletion strain that could not be rescued by addition of LiaX (**Figure 2D**). These findings indicated that MadRS ultimately controls the effector systems responsible for CAMP resistance in *E. faecalis*.

To test this hypothesis, we made targeted deletions of the genes encoding MadEFG and DltA in the OG117Δ*liaX* background, where the LiaFSR system is constitutively expressed due to loss of the regulatory C-terminal domain of LiaX. Consistent with previous results, the OG117Δ*liaX* strain showed enhanced survival in the presence of LL-37 as compared to OG117 (71.5% vs 8.31%, p=0.0001, **Figure 2E**). Deletion of *madEFG* in this background led to a decrease in survival (71.5% vs 33.7%, p=0.013), suggesting this system plays a significant role in membrane defense against LL-37, even in the presence of a constitutively active LiaFSR response. The loss of *dltA* led to a 27.9% reduction in survival after exposure to LL-37 in the OG117Δ*liaX* background that was not statistically significant. Interestingly, in the OG117Δ*liaX* background loss of *madEFG* led to a significant decrease in survival in the presence of HBD3 (70.9% vs 36.4%, p=0.013, **Figure 2F**), suggesting that the MadRS system can respond to CAMPs such as β-defensins and cathelicidins.

MadEFG did not appear to directly impact daptomycin susceptibility; however, a reduction in the MIC of daptomycin from 12 to 2 μg/mL was observed in the OG117Δ*liaX*Δ*dltA* strain, and the MIC was restored on complementation of *dltA* in the native chromosomal location. Since D-alanylation of teichoic acids has been implicated in changes of cell surface charge and daptomycin resistance in *S. aureus*, we evaluated cell surface charge using a cytochrome c binding assay (**Supplemental Figure 2**). As compared to OG117, OG117Δ*liaX* displayed a small but significant increase in positive cell surface charge. This increase was abolished in the OG117Δ*liaX*Δ*dltA* strain and restored on complementation of *dltA*. In the OG117Δ*dak* background, there was a small increase in cell surface charge of the *dak* deletion mutant, but no significant changes across any of the other strains. Thus, a simple electrostatic repulsion model cannot sufficiently explain the phenotypic changes observed in *E. faecalis*.

### Loss of *dak* alters the membrane lipid environment and leads to activation of MadRS in the absence of antibiotic stress

We have previously shown that *E. faecalis* with a truncated C-terminal Dak deletion have a growth defect and more rigid cell membrane.^7^ The *dak* gene is predicted to encode a fatty acid kinase which, in *S. aureus*, has been shown to mediate phosphorylation of exogenous fatty acids for use in membrane biosynthesis.^13^ In the absence of *dak*, the endogenous type II fatty acid synthesis pathway (FASII) supplies acyl-chains for phospholipid synthesis. Consistent with this hypothesis, OG117Δ*dak* has increased sensitivity to triclosan (4 µg/mL), an inhibitor of the FabI enoyl-ACP reductase of FASII, as compared to wild type OG117 or the *dak* complemented strain (both 16 µg/mL). During construction of the OG117Δ*dak* derivatives, we noted that OG117Δ*dakmadS_A202E_* and subsequent strains had a restoration of growth (**Supplemental figure 3**). Whole genome sequencing of these strains showed a serine to tyrosine change at amino acid 36 of FabT, a MarR transcriptional repressor which binds upstream of the Fab operon (encoding the FASII enzymes). Based on alignment with FabT from *Streptococcus pneumoniae*, for which the crystal structure has been solved, this change would be predicted to fall in the DNA binding domain, potentially impairing the binding of the repressor and, thus, resulting in constitutive expression of the Fab operon.^17^ Colonies with the mutations in FabT had no change in triclosan susceptibility compared to OG117Δ*dak* (4 µg/mL), consistent with continued reliance on endogenous FA synthesis.

To assess the membrane changes associated with the absence of *dak*, we performed hydrophilic interaction chromatography-mass spectrometry (HILIC-MS) to characterize the phospholipids of OG117, OG117Δ*dak*, and the OG117Δ*dak*::*dak* complemented strain, as well as OG117Δ*dakmadS_A202E_* (**Figure 3A and 3B**) during exponential growth and stationary phase. In exponential phase, OG117Δ*dak* had a significant decrease in the abundance of PG, Lysyl-PG (LPG), and diacyglycerol (DG) as compared to OG117. In both the *dak* complemented and the OG117Δ*dakmadS_A202E_* strain (with the mutation in *fabT*), there were no significant changes in phospholipid abundance. In stationary phase, the OG117Δ*dakmadS_A202E_* strain had a significantly higher abundance of PG and LPG as compared to OG117, consistent with the increased expression of *mprF2* (the product of which synthesizes LPG) in the *madS_A202E_* mutant. The acyl chain composition of each lipid species in exponential and stationary phase is shown in **Figure 3C** and **Supplemental Table 2**. Strains with a deletion of *dak* showed a relative increase in shorter chain and saturated fatty acids as compared to OG117 during exponential phase growth.

**Figure 3.**
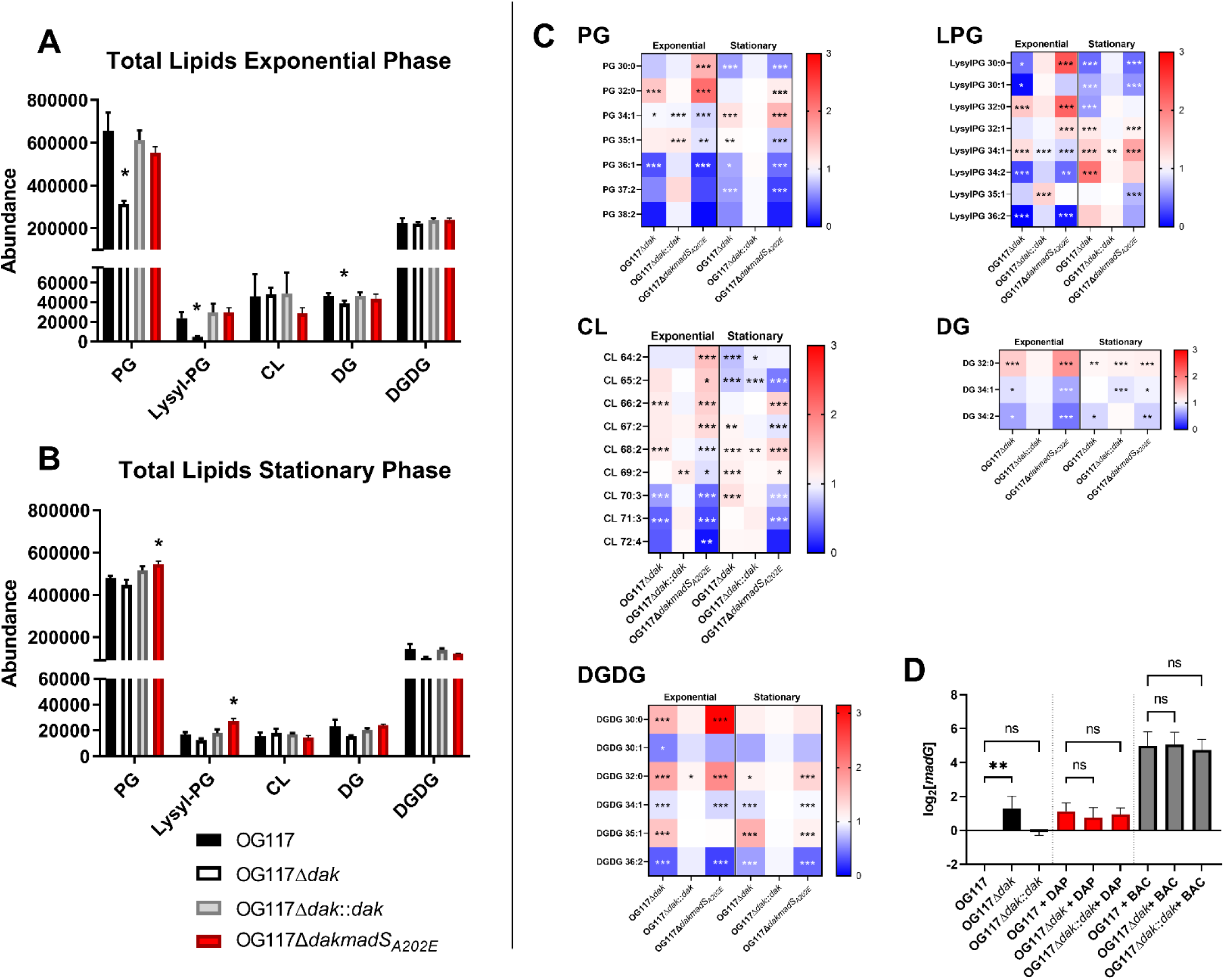
Deletion of *dak* is associated with changes in membrane fatty acyl composition and alteration of *madG* expression. Lipid analysis was carried out using Hydrophilic interaction chromatography-ion mobility-mass spectrometry (HILIC-IM-MS). Total lipids in the A) exponential growth and B) stationary phase were determined in the OG117 and *dak* deletion strains. Relative abundance is listed on the y-axis, and each major enterococcal lipid species is indicated on the x-axis. Significant differences in abundance of phosphatidylglycerol (PG), Lysyl-PG, and diacylglycerol (DG) were seen in the *dak* deletion strain at exponential phase, but not stationary phase. The OG117Δ*dakmadS_A202E_* strain developed a spontaneous mutation that led to de-repression of the *fab* operon (see text). No differences were observed in this strain in exponential phase, but there was a significant increase in PG and lysyl-PG at stationary phase. C) Fatty acyl chain composition of PG, lysyl-PG (LPG), cardiolipin (CL), DG, and digalactosyldiacylglycerols (DGDG) in exponential and stationary phase are shown as a heat map based on relative change from wild type OG117, with red representing an increase and blue representing a decrease in the relative proportion of each species. Carbon chain length and number of unsaturated bonds are given (e.g., PG 34:1). Both strains with the *dak* deletion showed a similar acyl chain profile, despite the constitutive activation of the FASII system in the OG117Δ*dakmadS_A202E_* mutant. D) Expression of *madG* at exponential phase in the *dak* deletion strains using qRT-PCR, without antibiotics (black bars), with daptomycin (DAP, red bars), and with bacitracin (BAC, grey bars). In the absence of antibiotic stress, there was a significant 2.5-fold increase in *madG* expression in the OG117Δ*dak* strain as compared to OG117. Under daptomycin and bacitracin stress, there was no significant difference in *madG* expression between the strains. *, p < 0.05; **, p < 0.01; ***, p < 0.001; ns, not significant.

To investigate whether the altered membrane environment associated with the deletion of *dak* influenced expression of the wild type MadRS system, we evaluated the expression of *madG* with and without antibiotic stress in the *dak* mutant strains during exponential growth (**Figure 3D**). In the absence of antibiotic stress, there was a 2-fold increase in the expression of *madG* in the *dak* deletion mutant as compared to OG1RF, similar to the 2-fold induction of *madG* expression seen after exposure to daptomycin in all strains. Bacitracin led to a robust expression of *madG* that did not differ across strains. Taken together, our results suggest that changes in phospholipid composition due to the deletion of *dak* both directly contribute to CAMP resistance and prime the transcriptional response of the MadRS system.

### MadRS is associated with CAMP dependent changes in survival of *Caenorhabditis elegans*

Using a *C. elegans* infection model, we examined the role of the MadRS system in protecting enterococci from CAMPs produced by the innate immune system. In wild type *C. elegans*, there was an attenuation of the OG1RFΔ*madR* strain as compared to OG1RF (median survival 11 days versus 6 days, p<0.0001, **Figure 4A**). This phenotype was reversed with complementation of *madR* on a plasmid with minimal differences between OG1RF and the complemented strain (median survival 6 versus 7 days, p=0.01). To examine if this survival difference was influenced by the presence of host CAMPs, we performed the same survival assay in *C. elegans* lacking the *pmk-1* gene, which encodes an ortholog of the p38 MAP kinase necessary for production of CAMPs. **Figure 4B** shows that there were no differences in survival between Δ*pmk-1* worms infected with OG1RF, OG1RFΔ*madR*, or the complemented strain, supporting the notion that the presence of CAMPs mediates survival of *C. elegans* infected with *E. faecalis* lacking MadR, rather than a general change in strain virulence.

**Figure 4.**
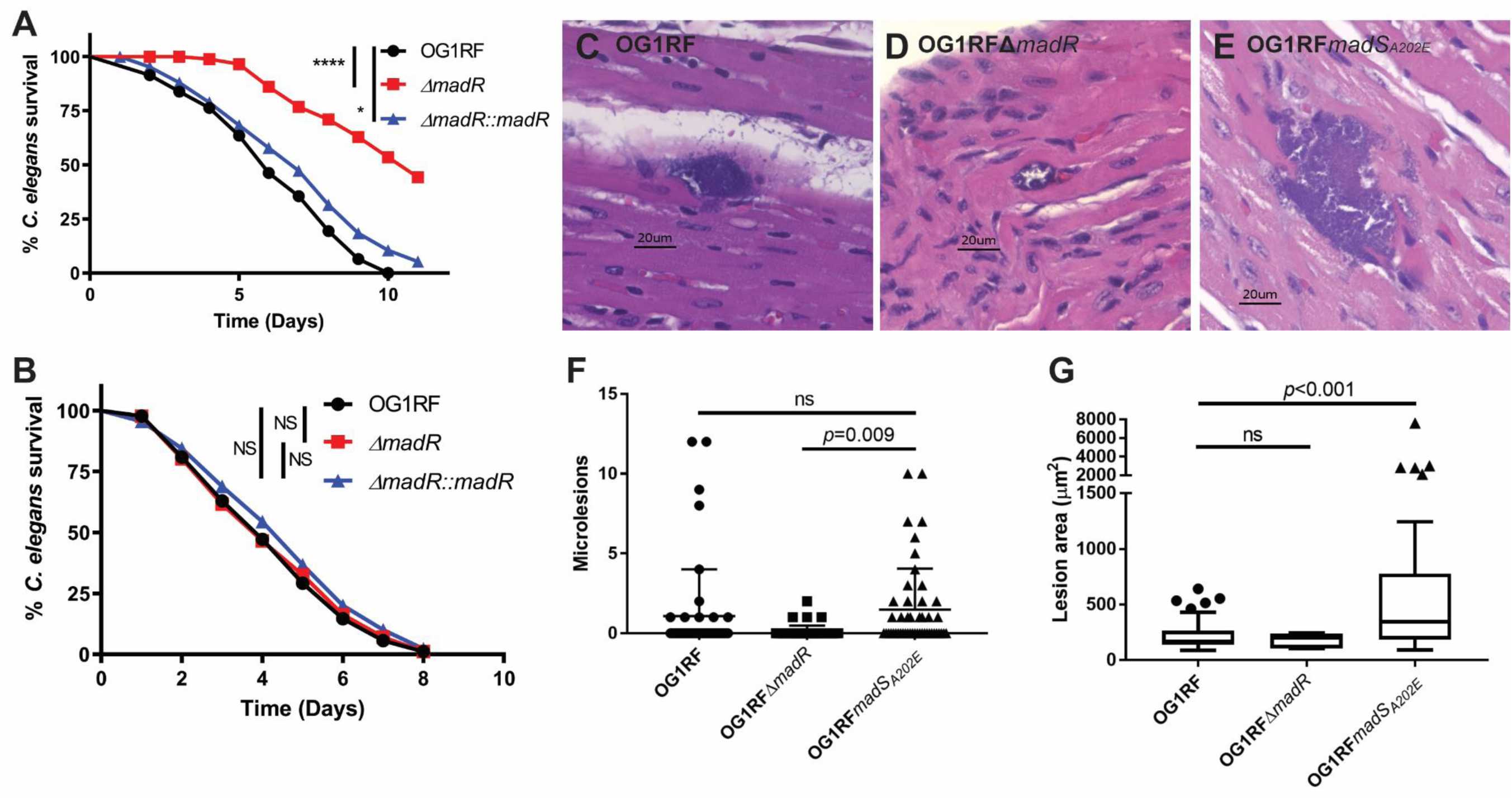
The MadRS system protects *E. faecalis* from the host immune response *in vivo*. Nematode survival in a C. elegans model of infection with A) wild type nematodes and B) *pmk-1* knockouts deficient in antimicrobial peptide production. Nematodes showed increased survival when infected with the OG1RFΔ*madR* strain as compared to the wild type OG1RF or when *madR* was complemented in trans on a plasmid, however, no differences in survival were seen in the nematodes unable to produce AMPs. C-D) Cardiac microlesions were assessed in a mouse model of peritonitis for the OG1RF, OG1RFΔ*madR*, and OG1RF*madS_A202E_* strains. Murine hearts were formalin fixed and embedded in paraffin, bisected, then sectioned on a microtome. Representative micrographs of hematoxylin and eosin stained sections are shown. F) Average number of cardiac microlesions per section. Eight sections were evaluated per animal and 48 in total for each strain. Differences in the mean number of microlesions between each strain were determined using one-way ANOVA. P-values are indicated for each comparison on the graph. G) Lesion area in µm^2^ are given as box and whisker plots of inner quartile range, with error bars plotted by the method of Tukey. Outliers are shown as individual data points. Statistical differences between strains were assessed using the Kruskal-Wallis test. P-values are indicated for each comparison on the graph. ns, not significant.

### The *madS_A202E_* allele is associated with an increased burden of cardiac microlesions in a mouse peritonitis model of infection

Given the importance of host CAMPs to the innate immune response, we sought to investigate the *in vivo* impact of the MadRS system in a mouse model of peritonitis. We examined overall survival, bacterial burden in the spleen, and the average number and size of cardiac microlesions as assessed by histopathology of cardiac tissue from infected mice.^18^ There was no difference in median survival or bacterial burden in the spleen between OG1RF, OG1RFΔ*madR*, and OG1RF*madS_A202E_*(**Supplemental Figure 5**). However, mice infected with OG1RF*madS_A202E_* showed an increase in the mean number of cardiac microlesions per histopathological section as compared to OG1RFΔ*madR* (1.48 vs. 0.1, p=0.009, **Figure 4C-G**), and a significantly larger lesion area as compared to OG1RF (p<0.001). The findings suggest that the MadRS system may play a critical role in the immune mediated clearance of deep seated cardiac enterococcal infections.

## Discussion

The potential for cross resistance between daptomycin and CAMPs of the innate immune system is of particular concern, since enterococcal exposure to CAMP in the GI tract has the potential to prime antibiotic resistance and serve as a precursor to mucosal translocation and bloodstream infections, particularly in the immunocompromised host.^19–21^ In this study, we characterized a specific defense network that protects the enterococcal cell envelope and dissected the effector genes responsible for resistance to CAMPs and peptide-like antibiotics.

Using transcriptional analyses, we identified the regulatory network of the MadRS system, showing that it has a much broader cell envelope protective effect than previously known (**Figure 5**). Consistent with prior results, MadR was responsible for the expression of MadLM and that activation of the system also resulted in an increase in the expression of the *dlt* operon.^22^ In addition, MadR appears to regulate the expression of the peptidoglycan hydrolase SalA, the lysylphosphatidylglycerol synthetase MprF2, and the novel *madEFG* operon, which had not been previously characterized in *E. faecalis*. The *madEFG* genes share sequence identity with the *spr0693-0695* operon in *Streptococcus pneumoniae*, which has been shown to encode a MacAB-like efflux pump with LL-37 as a potential substrate.^23^ The product of *spr0693* was predicted to form a hexameric channel of ∼155 Å, potentially spanning the peptidoglycan layer. Thus, we postulate that MadEFG is likely to function by removing CAMPs at the membrane interface and transporting them past the peptidoglycan layer. The other components of the MadRS regulon alter the cell envelope by D-alanylation of wall teichoic acids via Dlt or production of lysyl-phosphatidylglycerol via MprF to prevent diffusion of CAMPs back to the membrane surface. Interestingly, no obvious homologues of *madEFG* are present in the genome of *Enterococcus faecium* DO, suggesting that there are important differences in the mechanism underlying membrane defense between the two species.

**Figure 5.**
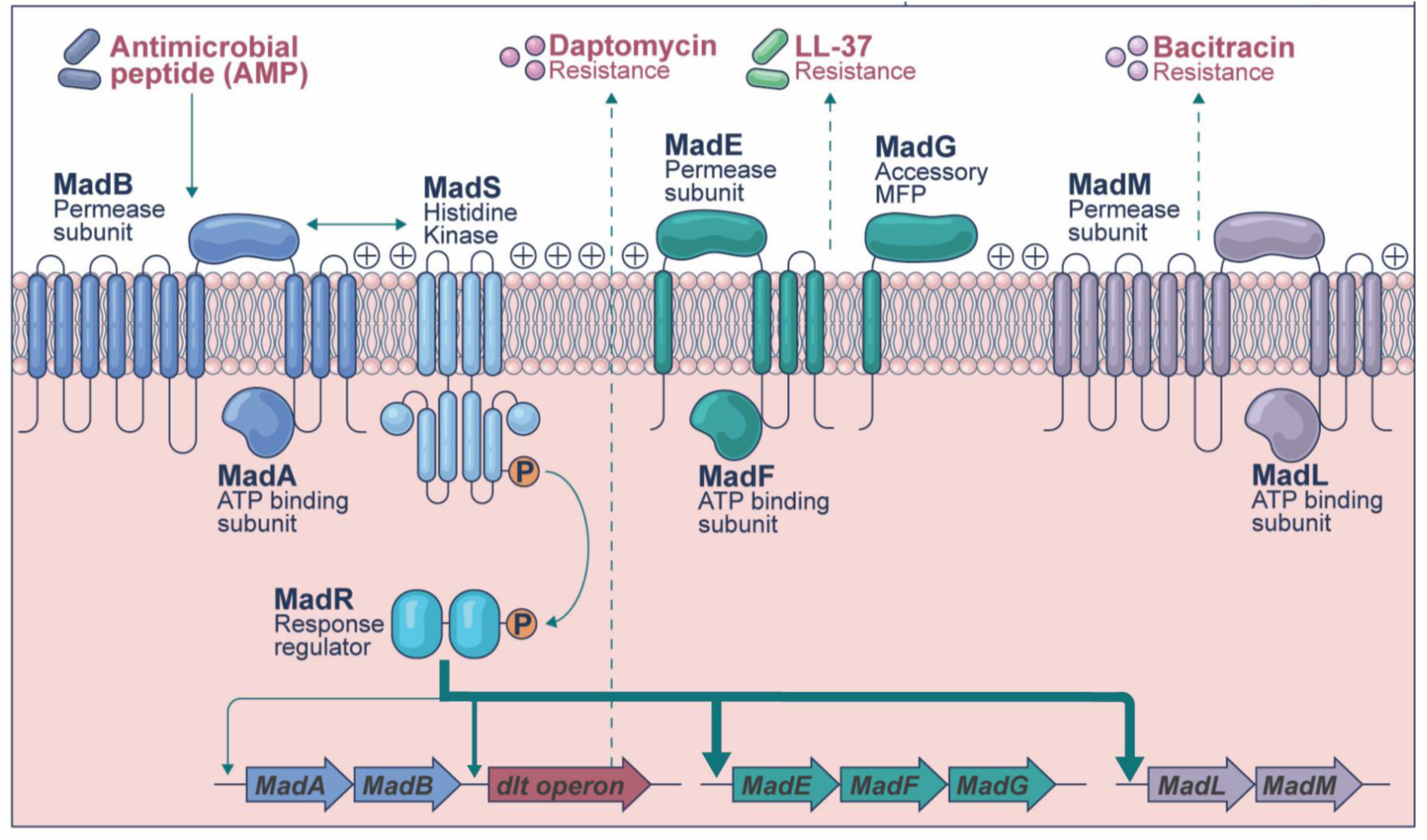
Model for the function of the Membrane Antimicrobial Peptide Defense System. The MadRS system controls a network of effector proteins that provide a coordinated and specific defense against CAMPs and CAMP-like antibiotics. The MadS sensor histidine kinase is activated by the flux-sensing mechanism of the constitutively expressed MadAB ATP-binding cassette (ABC) transporter in the presence of AMPs. This leads to phosphorylation of the MadR response regulator and upregulation of the genes encoding MadEFG, MadLM, and the *dlt* operon, as well as *mprF2* and the gene encoding a putative peptidoglycan hydrolase SalA (not pictured). These targets each provide protection from their specific substrates including bacitracin (MadLM), LL-37 and HBD3 (MadEFG), and daptomycin (DltABCD). After removal of CAMPs from the membrane by the ABC transporters, D-alanylation of teichoic acids and lysylation of phosphotidylglycerol alters cell surface charge and prevents diffusion of these molecules back to the cell membrane surface. Alterations of the cell membrane lipids (not pictured) via loss of function in *dak*, or other mechanisms, primes expression of the MadRS system and may also directly interfere with CAMP binding.

Alterations of the membrane bilayer, as we observed in the *dak* knockout, may provide a further barrier to effective CAMP action. Loss of Dak function resulted in a membrane lipid fatty acid profile that contained significantly more saturated fatty acids, in part explaining our prior observation of decreased membrane fluidity (presumably from a tighter lipid packing order) in the Dak mutant.^7^ These changes were sufficient to decrease the activity of HBD3, and also appear to prime the MadRS system by leading to an increase in the baseline transcription of target genes, thus, preparing the cell to defend against CAMPs. Thus, the membrane lipid environment has both a direct and indirect role in antimicrobial resistance, and these findings may explain why mutations in both phospholipid metabolism and cell envelope stress response networks emerge together in daptomycin-resistant isolates.

Importantly, the MadRS response appears to complement the global stress response of LiaFSR with a targeted set of effector proteins. Recently, a link between the LiaFSR and MadRS systems was demonstrated in *E. faecalis*, with activation of the LiaFSR system associated with increased expression of MadRS.^24^ Similar to the Bce-type transporters, MadAB-MadS are likely to function via a flux-sensing mechanism, where the transport activity of MadAB is coupled to the kinase activity of MadS.^25^ The data presented here suggest that the emergence of daptomycin resistance via MadRS arises via a decoupling of the MadS kinase and the MadAB flux sensor, as the MadS_A202E_ mutant showed increased expression of the genes of the MadR regulon, even in the absence of antibiotic stress. This decoupling bypassed the potential transcriptional regulation of the MadRS system by LiaFSR, and activation of MadRS results in a survival advantage against CAMPs, even in the absence of *liaR*.

Moreover, the results from the *C. elegans* and mouse peritonitis models suggest that the MadRS system also plays a major role against CAMPs *in vivo*, and resistance arising via this pathway may have implications for the clearance of deep-seated infections such as infective endocarditis. Interestingly, we did not see a difference in bacterial burden in the spleen. One possibility is that immune mediated clearance via macrophages and neutrophils may be preserved via activity of alternate peptides such as HNP-1, the activity of which was not affected in our assay conditions. Since peptides such as β-defensins and cathelicidins are an important part of mucosal barrier defense, this resistance pattern may predispose to invasive infection and bacteremia, particularly in immunocompromised hosts with neutropenia.^26,27^ Further work is needed to understand the role of this system with respect to colonization of the GI tract and subsequent risk for translocation and infection.

In summary, the MadRS system controls a network of genes in *E. faecalis* that are capable of conferring resistance to the antibiotic daptomycin and host immune peptides including HBD3 and LL-37. In *E. faecalis*, survival in the presence of these CAMPs is dependent on the presence of *madEFG*, a three gene operon predicted to encode a MacAB-like efflux pump, with alterations in membrane lipids also contributing to HBD3 resistance. Mutations leading to constitutive activation of the MadRS system are able to provide daptomycin resistance independent of LiaFSR, and our *in vivo* data suggest this system may play an important role in the persistence of enterococcal infections.

## Acknowledgements

The authors wish to thank Rafael Rios for assistance with the RNA-Seq analysis. WRM is supported by a National Institute of Health, Institute of Allergy and Infectious Diseases (NIH/NIAID) grant K08 AI135093. CAA is supported by a NIH/NIAID K24 AI121296, R01 AI148342, R01 AI134637, and P01 AI152999. DAG is supported by R01 DE027608, R01 AI150045, and R21 AI167124. SLE was supported by a training fellowship from the Gulf Coast Consortia, Texas Medical Center Training Program in Antimicrobial Resistance (TPAMR) (NIAID) T32 AI141349. YS is supported by NIAID R01 A1080714. LX is supported by NIH/NIAID R01 AI136979.

## Conflict of Interest

WRM has received grant support from Merck and royalties from UpToDate. CAA has received royalties from UpToDate. All other authors have no conflicts to disclose.

## Data Availability Statement

Datasets for the whole genome sequencing and RNA-seq are available at the National Center for Biotechnology Information (NCBI) under Bioproject Accession number PRJNA1025297.

## Online Methods

### Bacterial strains, plasmids and growth conditions

The bacterial strains and plasmids used in this work are listed in Supplemental Table 3. All enterococcal strains were grown on brain heart infusion (BHI, Becton Dickinson) agar or in BHI broth on rotating incubator at 37° C unless otherwise specified. The *Escherichia coli* strains EC1000 and TG-1 were used for propagation of the plasmids pHOU1 and pAT392, respectively and grown on Luria-Bertani (LB, Becton Dickinson) agar or in broth at 37° C supplemented with 25 µg/mL gentamicin (Sigma). Minimum inhibitory concentrations (MIC) were performed by gradient diffusion test on Mueller-Hinton agar (MH, Becton Dickinson) for daptomycin, ampicillin, vancomycin (Etest, bioMerieux), and bacitracin (MTS, Liofilchem). MICs for triclosan were performed by broth microdilution in MH broth using two-fold serial dilutions of triclosan (range 0.125-128 µg/mL, Sigma). Bacterial growth curves were performed in a 96 well plate (Fisher) with inoculation of approximately 5x10^5^ CFU/mL final concentration into a volume of 200 µL BHI. Optical density at 600 nm was measured on a Synergy H1 plate reader (BioTek) over 24 hours at 15-minute intervals with 30 seconds of orbital shaking prior to each read.

### Genetic manipulation of strains

Chromosomal deletions were generated using a derivative of OG1RF with the *E. faecalis* ATCC 4200 CRISPR1 *cas9* gene inserted in a neutral genomic insertion site to aid in genetic manipulation.^28,29^ A guide RNA for directed cleavage of the target sequence was provided in trans in pCE under a constitutive pBac promoter. The pCE plasmid also contained a crossover region containing approximately 800 bp up and downstream of the gene or region of interest.

PCR fragments were amplified from empty pCE, or purified enterococcal genomic DNA templates, with approximately 20 bp complementary overhangs and assembled via Gibson Assembly (Gibson Assembly Master Mix, New England Biolabs) by incubation in a thermal cycler at 50° C for 1 hour. Primers used for each construct can be found in Supplemental Table 4. After assembly, the reaction was diluted 1:3 in nuclease free water and 6 µL was transformed into chemically competent *E. coli* DH5α (High Efficiency, New England Biolabs). Transformants were selected on LB agar with 15 µg/mL of chloramphenicol and screened with primers to ensure correct plasmid and insert. Overnight cultures of *E. coli* with the construct of interest were grown in LB broth with 15 µg/mL of chloramphenicol, plasmid was isolated (Wizard Plus SV Miniprep DNA Purification System, Promega), and then electroporated into electrocompetent *E. faecalis* OG117 or derivative as needed.^30^ Transformants were selected on BHI agar with 15 µg/mL of chloramphenicol. For constructs that could not be directly electroporated, they were first electroporated into *E. faecalis* CK111, then filter mating with the final strain was performed as previously described.^31^ Transconjugants were selected on BHI plus 15 µg/mL of chloramphenicol and 25 µg/mL fusidic acid. Colonies were then screened by PCR to ensure the presence of the correct gene knock out or complementation. Curing of the pCE plasmid was performed by streaking colonies on MM9-YEG plates containing p-chloro-phenylalanine for counterselection. Colonies that grew on fusidic acid, but not chloramphenicol or spectinomycin, were screened for presence of the plasmid and region of interest by PCR. All constructs were verified by sanger sequencing over the entire crossover region, and whole genome sequencing.

### RNA sequence analysis and qRT-PCR

RNA-seq analysis was performed for the strains OG1RF, OG1RFΔ*madR*, and OG1RF*madS_A202E_* as previously described.^10^ Briefly, strains were grown overnight in tryptic soy broth (TSB, Becton Dickinson) plus 50 mg/L of calcium chloride, then diluted into fresh TSB with Ca^2+^ alone or with daptomycin at ½ of the MIC and grown to mid-exponential phase (OD_600nm_ 0.5-0.7) with agitation at 37° C. Cell pellets were harvested by centrifugation of 2 mL of mid-exponential culture and stored at −80° C until extraction. Three independent biological replicates were performed for each strain and treatment condition. Total RNA was obtained with the Ribopure RNA purification kit (Ambion, Life Technologies) followed by DNase treatment, and subsequent analysis for quality using Nanodrop (Thermo Scientific) and Bioanalyzer (Agilent).

Depletion of ribosomal RNA was performed using the Ribozero kit (Illumina). cDNA libraries were prepared with the TrueSeq RNA Sample Preparation System v.2 (Illumina), and samples were sequenced on an Illumina HiSeq2000. Reads were mapped using the OG1RF genome, using the uninduced OG1RF as a reference for baseline expression. Reads per kilobase of gene per million reads (RPKM) was calculated with EDGE-pro v1.3.1.^32^ An adjusted p-value threshold of p < 0.01 using the Benjamini-Hochberg procedure to control the false discovery rate was used to identify the set of differentially expressed genes for each comparison with DESeq2.^33^

Quantitative real-time PCR was performed as previously described.^10^ Cells were grown as above, in antibiotic induced and uninduced conditions, with either daptomycin or bacitracin-zinc at ½ the respective MIC. Total RNA was isolated using the Pure Link RNA Isolation kit (Invitrogen), genomic DNA was removed with the Turbo DNase kit (Ambion), and cDNA synthesis was performed using SuperScript II Reverse Transcriptase (Invitrogen). Expression of *madG*, *madL*, *madA*, and *dltA* were evaluated by qRT-PCR using SYBR Green in a CFX96 Touch Real-Time PCR Detection System (Bio-Rad). Expression level was quantified relative to the control strain, OG1RF, without antibiotic exposure. Values were normalized to the expression of *gyrB* as a housekeeping gene. Primers used for qRT-PCR are listed in Supplemental Table 4. Fold-change was calculated using the Pfaffl method and primer efficiency was determined for each reaction with the LinRegPCR program. Differences in gene expression were determined using two-way ANOVA with Tukey’s test for multiple comparisons, with a p-value of < 0.05 (Graph Pad Prism). Each experiment was performed in biological and technical triplicates.

### Antimicrobial peptide killing assays

Antimicrobial peptide killing assays for LL-37 (50 µg/mL, Anaspec, Cat. #AS-61302), HBD3 (6 µg/mL, Anaspec, Cat. #AS-60741), and HNP-1 (10 µg/mL, Anaspec, Cat. #AS-60743) were performed as previously described with modifications as follows.^34,35^ All strains to be tested were grown overnight in 5 mL of BHI broth on a rotary incubator at 37° C. Overnight cultures were diluted 1:100 in 5 mL of fresh BHI broth and grown to mid-exponential phase (OD_600nm_ 0.5-0.7). Strains were then adjusted to 0.5 McFarland (OD_600nm_ 0.08-0.1) in the respective assay buffer: RPMI + 5% LB for LL-37 and HBD3 (HyClone RPMI 1640 without glutamine and without phenol red, VWR), and 10 mM KH_2_PO_4_, pH 7.4 for HNP-1. Each respective McFarland standard was further diluted in assay buffer to a final inoculum of 1x10^4^ CFU/mL and 5 µL of this suspension was added to 45 µL of assay buffer with and without each antimicrobial peptide in technical triplicate. Cells were then incubated for 2 hours at 37°C. After incubation, two serial 10-fold dilutions were made and 25 µL of each replicate was plated on fresh BHI agar plates. Spots were allowed to dry, then incubated overnight at 37° C for colony counts. Percent survival for each strain was calculated as CFU for peptide treated sample divided by the CFU for the growth control sample times 100. Each assay was performed in three independent biological replicates for each strain. Differences in mean survival were determined using one-way ANOVA with Dunnet’s multiple comparisons test, with a significance threshold of p < 0.05 (GraphPad Prism).

### Cytochrome c assay

Cell surface charge was determined by cytochrome c assay.^36^ Overnight cultures were grown in BHI broth to stationary phase, then pelleted by centrifugation, and resuspended in 1 mL of 20 mM MOPS buffer, pH 7.0 (Sigma). After resuspension, the suspension was transferred to a microcentrifuge tube and washed twice with MOPS buffer. Washed cells were adjusted to a final density of OD_600nm_ of 7.0, and 0.25 mg/mL of cytochrome c (bovine heart, Sigma) was added to each sample and a cell free MOPS buffer control. Cytochrome c was allowed to bind for 15 minutes at 37° C with shaking, then cells were centrifuged at 14,000 relative centrifugal force for 5 minutes. An aliquot of 100 µL times 3 technical replicates was pipetted from the supernatant, with care not to disturb the cell pellet, and transferred to a clear plastic 96-well plate.

Absorbance of each sample was read at 410 nm on a spectrophotometer (Synergy H1, BioTek). The percentage bound cytochrome c was determined by calculating the difference from the absorbance of the cytochrome c cell free control. Statistical comparison in the difference between the mean level of binding was performed using one-way ANOVA with Dunnett’s test for multiple comparisons, with a significance threshold of p < 0.05 (GraphPad Prism).

### Membrane lipid extraction

Lipids were extracted by the method of Bligh and Dyer, and as previously described.^37–39^ Briefly, bacteria were grown into mid-exponential phase or stationary phase, respectively, washed in 1x phosphate buffered saline (PBS), then pelleted and dried in a centrifuge under vacuum. The dried pellets were resuspended in 1 mL of sterile water (LC grade, Thermo Fisher Scientific), then sonicated in an ice bath for 30 minutes to homogenize the sample. A 4 mL chilled solution of chloroform and methanol (1:2 v/v, LC grade, Thermo Fisher Scientific) was added to each tube and vortexed for 5 minutes. An additional 1 mL chilled chloroform and 1 mL chilled water were added to each sample, vortexed for 1 minute, then centrifuged at 2,000 x *g* for 10 minutes at 4° C. The organic layer was collected to a 10 mL glass centrifuge tube (Fisher Scientific) and dried in a vacuum concentrator. The dried lipids were reconstituted in 500 µL of a 1:1 mixture of chloroform and methanol and stored at −80° C until further use.

### Determination of membrane lipids by hydrophilic interaction chromatography-ion mobility-mass spectrometry (HILIC-IM-MS)

A 5 µL aliquot of each lipid extract was placed in LC vials, dried under nitrogen, and redissolved in 100 µL of a 2:1 mixture of acetonitrile and methanol (LC grade, Thermo Fisher Scientific). The lipids were separated on a Waters UPLC (Waters Corp., Milford, MA, USA) using a Phenomenex Kinetex HILIC column (2.1x100 mm, 1.7 µm) at a temperature of 40° C and a flow rate of 0.5 mL/minute.^40,41^ A sample injection volume of 5 µL was used for each sample. The solvent systems consisted of (A) 50% acetonitrile/50% water with 5 mM ammonium acetate (Optima LC-MS grade, Thermo Fisher Scientific) and (B) 95% acetonitrile/5% water with 5 mM ammonium acetate. The solvent gradient was 100% B 0-1 minutes, 90% B 4 minutes, 70% B 7-8 minutes, 100% B 9-12 minutes.

Lipid analysis was performed on a Waters Synapt G2-XS platform. Samples separated by the UPLC were injected via the electrospray ionization (ESI) source. ESI capillary voltages of +2.0 were used for positive analyses and −2.0 kV for negative analyses. Additional ESI conditions utilized were a sampling cone, 40 V; extraction cone, 80 V; source temperature, 150 °C; desolvation temperature, 500 °C; cone gas, 10 L/hr; desolvation gas, 1000 L/hr. Mass calibration over *m/z* 50-1200 was performed with sodium formate. Calibration of ion mobility (IM) measurements was performed as previously described.^42^ IM separation was performed with a traveling wave height of 40 V and velocity of 500 m/s. Data was acquired for *m/z* 50-1200 with a 1 sec scan time. Untargeted MS/MS (MS^E^) was performed in the transfer region with a collision energy ramp of 35-45 eV. Mass and drift time correction was performed post-acquisition using the leucine enkephalin lockspray signal.

Data analysis was performed used the Progenesis QI software (Nonlinear Dynamics, Milford, MA, USA) for alignment, peak detection, and normalization. A pooled quality control sample was used as the alignment reference. All PC and PE lipid standards used for CCS calibration were purchased from Avanti Polar Lipids.^43^ The default “All Compounds” method of normalization was used to correct for variation in the total ion current amongst samples. Lipid identification was based on *m/z* (within 10 ppm mass accuracy), retention time, and CCS with an in-house version of LipidPioneer, modified to contain the major lipid species observed in *E. faecalis*, including DGs, DGDGs, PGs, CLs, and LysylPGs with fatty acyl compositions ranging from 25:0 to 38:0 (total carbons : total degree unsaturation), and LiPydomics.^41,44,45^

### Caenorhabditis elegans *infection model*

A *C. elegans* infection model was used to assess the impact of the loss of the MadR response regulator on survival in wild type and *pmk-1* deficient nematodes.^46,47^ Synchronized young adult nematodes were infected with OG1RF, OG1RFΔ*madR*, or OG1RFΔ*madR* complemented with *madR* in trans on pAT392 on BHI agar supplemented with 10 µg/mL of gentamicin.

Approximately 60-90 nematodes were transferred to a liquid media of 80% MW9 buffer and 20% BHI broth in 96 well plates. Plates were incubated at 25° C and evaluated daily for nematode survival, which was assessed via movement on stimulation with a sterile platinum wire probe. Survival curves were plotted using Kaplan-Meier log rank analysis, a difference in median survival from three independent runs was considered significant with a p-value of < 0.05.

### Mouse peritonitis infection model

A previously established mouse peritonitis model was used for this study.^48,49^ The protocol was approved by the University of Texas Health Science Center (UTHSC) Animal Welfare Committee (AWC-19-0058). The lethal dose for 50% of animals (LD_50_) for each strain was calculated by the method of Reed and Muench.^50^ Briefly, 4-6 week old female outbred ICR mice (Envigo) weighing approximately 25 grams were inoculated (n=6 per strain) via intraperitoneal injection of approximately 5x10^8^ CFU/mL in 50% sterile rat fecal extract. Mice were followed for 96 hours post infection, death was assessed approximately every 6 hours through the first 48 hours, then once daily through the end of the experiment. Differences in survival were determined by Mantel-Cox (log-rank) test. At the time of death, spleen and heart were aseptically excised. Spleens were added to pre-weighed bags, total tissue mass was calculated, then the organs were homogenized with a paddle blender (Stomacher 80 Biomaster, Seward Laboratory Systems). A total of 3 mL of sterile saline was added to each bag and mixed by pipetting, then serial ten-fold dilutions were prepared and 50 µL of each dilution was plated in triplicate onto fresh BHI agar plates. Plates were incubated overnight at 37° C, colonies were enumerated, and normalized to the tissue mass. To assess for cardiac microlesions, hearts were placed in formalin at the time of removal then processed by the UTHSC pathology core laboratory.^18^ The organs were embedded in paraffin, bisected, then sectioned longitudinally on a microtome, with 32 sections per block. Every 4^th^ section was stained with hematoxylin and eosin (8 sections per animal). Microscopy was carried out on a Keyence BZ-X700 microscope. Each lesion was imaged, the number of microlesions was determined for each section, and differences between the mean number of microlesions per section for each strain were calculated using one-way ANOVA with Tukey’s test for multiple comparisons and a significance threshold of p < 0.05. The area of each lesion was determined using FIJI software and differences between the strains were determined using Kruskal-Wallis test with Dunn’s test for multiple comparisons.^51^

## Notes

### Summary of Updates

Supplemental files uploaded.

