## Supplemental Fig 1-5, Supplemental Table 2-4 for "Membrane Lipids Augment Cell Envelope Stress Signaling and Resistance to Antibiotics and Antimicrobial Peptides in *Enterococcus faecalis*"

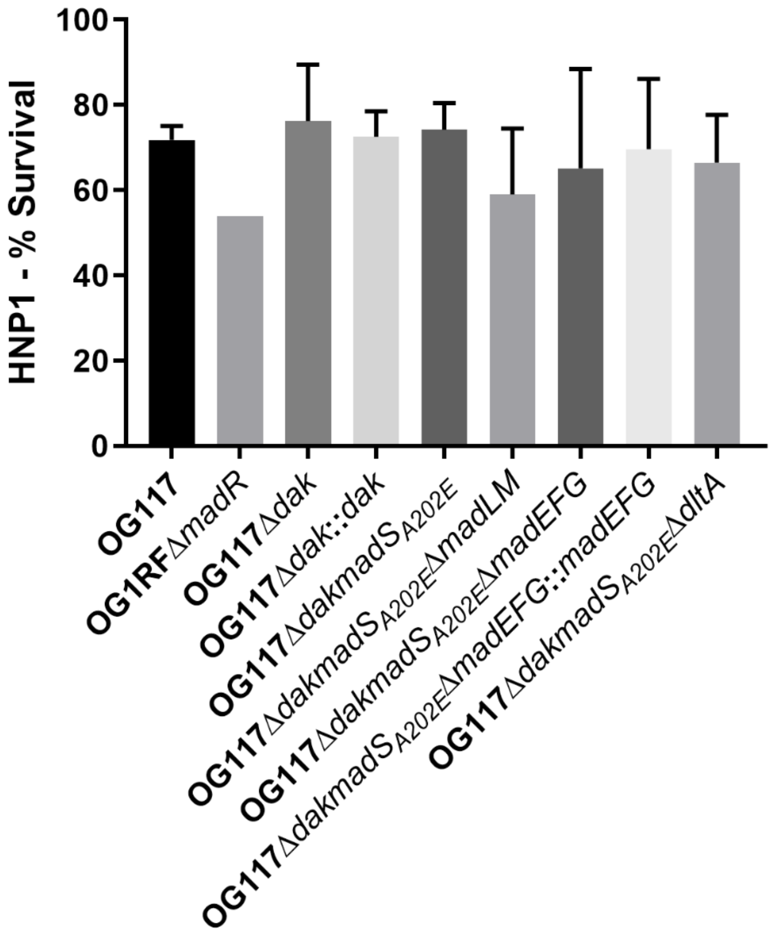


**Supplemental Figure 1. HNP1 peptide killing assay.** Bacterial isolates were incubated with HNP1 at 10 µg/mL. Percent survival was calculated by dividing the number of colony forming units per milliliter (CFU/mL) after HNP1 exposure by the CFU/mL of assay buffer growth control. No significant differences were seen across any of the strains at the tested conditions, error bars represent standard deviation of three independent runs.

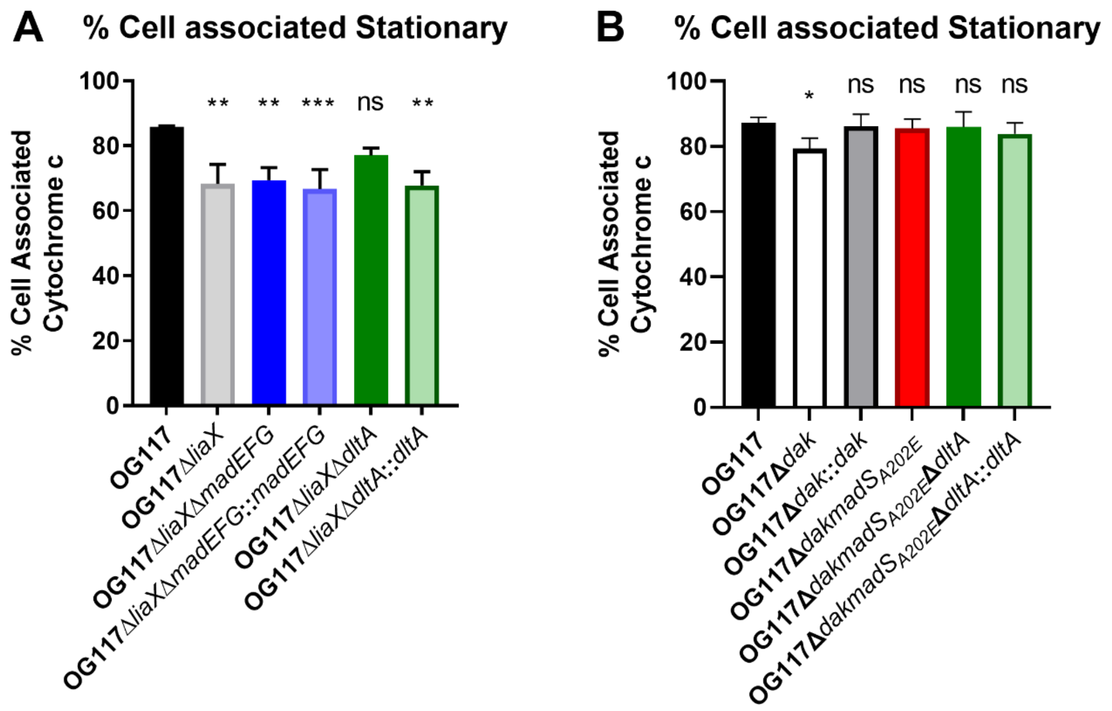


**Supplemental Figure 2. Cytochrome c binding assay.** Cell surface charge was assessed via binding of the cationic protein cytochrome c to cells in stationary phase. Increased positive cell surface charge (i.e., via Dlt mediated D-alanylation of lipo- and wall-teichoic acids) would decrease cell associated cytochrome c. **(A)** A decrease in cell associated cytochrome c was seen in the OG117Δ*liaX* background as compared to OG117. This difference in surface was abolished on deletion of *dltA* and restored with complementation of *dltA* in the native chromosomal location. **(B)** Deletion of *dak* was associated a significant decrease in cell associated cytochrome c as compared to OG117, however no significant changes were seen in the OG117Δ*dakmadS*$_{A202E}$ background. *, $p<0.05$; **, $p<0.01$; ***, $p<0.001$; ns, not significant. Error bars represent standard deviation of three independent runs.

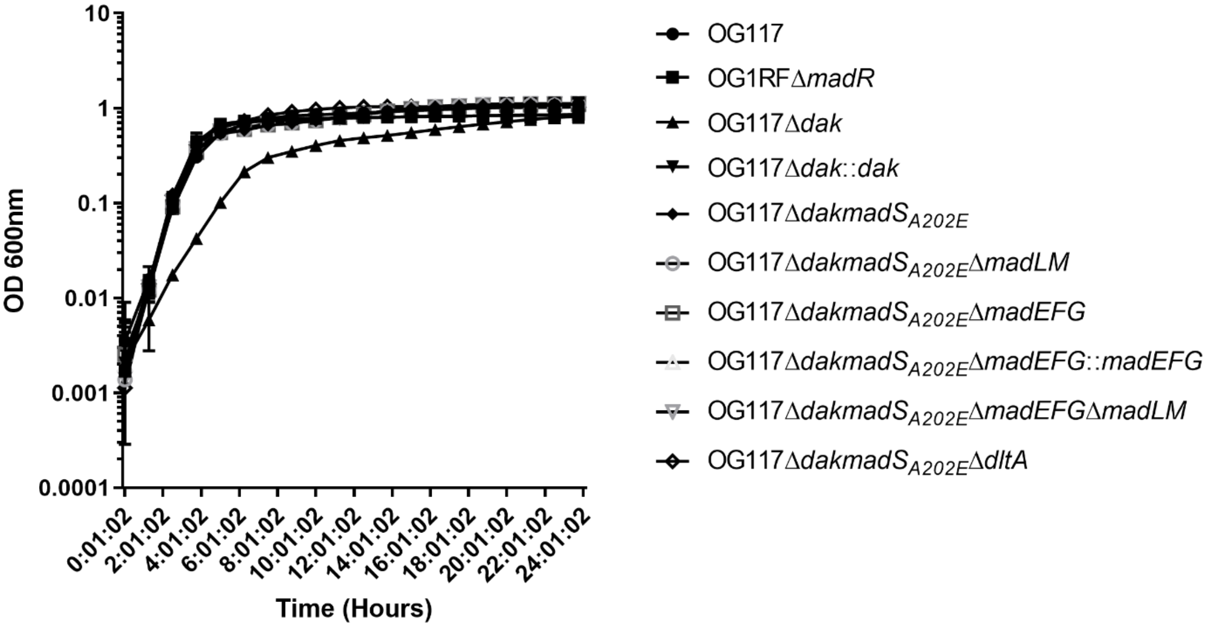


**Supplemental Figure 3. Growth curves for selected strains.** Bacteria were inoculated at approximately 1x$10^6$ CFU/mL and grown in brain heart infusion broth for 24 hours at 37° C in 96 well plate format. Optical density measurements at 600 nm were taken every 15 minutes, every $4^{th}$ measurement was graphed for clarity. Error bars represent standard deviation of at least 11 replicate wells.

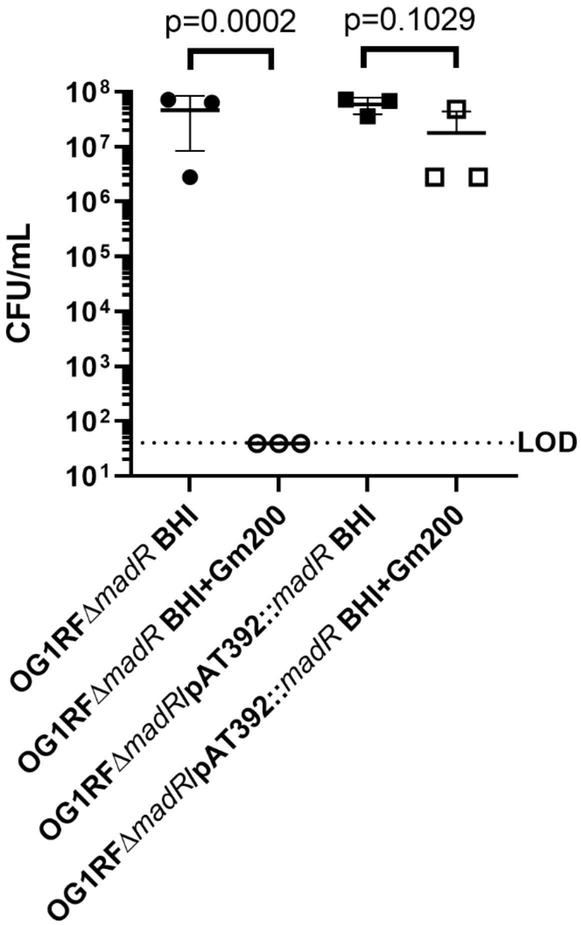


**Supplemental Figure 4. Plasmid stability for pAT392 in the absence of gentamicin exposure.** The stability of OG1RFΔ*madR* containing the plasmid pAT392::*madR* was determined after overnight passage in antibiotic free BHI media. Ten-fold serial dilutions for each strain were plated on BHI agar and BHI agar containing 200 µg/mL of gentamicin (Gm) for CFU determination. Statistical differences were determined by unpaired t-test of the log-transformed CFU. The limit of detection for the assay was 40 CFU/mL. The assay was performed in triplicate.

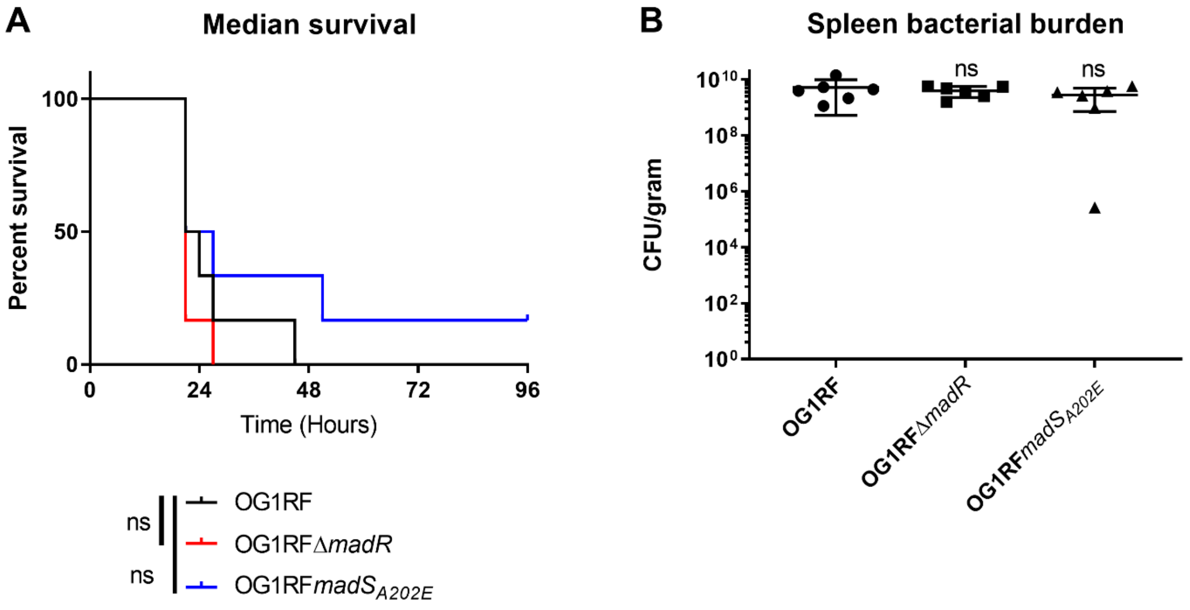


**Supplemental Figure 5.** ***E. faeacalis*** **mouse peritonitis model.** Female outbred ICR mice (n=6 per strain) were inoculated via intraperitoneal injection with approximately $5x10^8$ CFU/mL of bacteria in sterile rat fecal extract. **(A)** Kaplan-Meier survival curves for mice. Statistical differences in median survival were assessed by the log-rank test, there were no differences in median survival between the strains. **(B)** Bacterial burden from the spleen for each strain. Statistical differences were assessed by one-way ANOVA with Tukey's test for multiple comparisons. There were no significant differences between the strains.

| Exponential Phase | OG117 | OG117Δ*dak* | OG117Δ*dak*::*dak* |
|---|---|---|---|
| Phosphatidylglycerol | | | |
| PG 30:0 | 6.07 ± 0.36 | 4.71 ± 0.09 | 5.67 ± 0.16 |
| PG 32:0 | 11.36 ± 0.88 | **16.42 ± 1.34** | 11.73 ± 0.94 |
| PG 34:1 | 59.10 ± 1.55 | **57.36 ± 1.76** | **56.55 ± 1.73** |
| PG 35:1 | 17.00 ± 0.67 | **19.23 ± 0.69** | **19.66 ± 0.99** |
| PG 36:1 | 4.91 ± 0.21 | **1.53 ± 0.07** | 4.42 ± 0.2 |
| PG 37:2 | 1.39 ± 0.06 | 0.73 ± 0.03 | 1.79 ± 0.1 |
| PG 38:2 | 0.19 ± 0.02 | 0.018 ± 0.005 | 0.18 ± 0.013 |
| Lysylphosphatidylglycerol | | | |
| LysylPG 30:0 | 3.91 ± 0.7 | **1.71 ± 0.29** | 4.58 ± 0.65 |
| LysylPG 30:1 | 2.12 ± 0.13 | **0.08 ± 0.11** | 2.22 ± 0.2 |
| LysylPG 32:0 | 6.97 ± 1.48 | **9.98 ± 0.35** | 8.28 ± 1.63 |
| LysylPG 32:1 | 13.58 ± 0.82 | 12.19 ± 1.12 | 13.12 ± 1.17 |
| LysylPG 34:1 | 53.94 ± 1.54 | **67.67 ± 2.08** | **50.96 ± 1.64** |
| LysylPG 34:2 | 4.96 ± 0.57 | **1.25 ± 0.29** | 3.82 ± 0.5 |
| LysylPG 35:1 | 8.79 ± 1.29 | 7.12 ± 0.71 | **11.76 ± 1.65** |
| LysylPG 36:2 | 6.11 ± 0.53 | **0.005 ± 0.01** | 5.27 ± 0.45 |
| Cardiolipin | | | |
| CL 64:2 | 10.02 ± 1.53 | 9.12 ± 0.70 | 9.14 ± 0.35 |
| CL 65:2 | 3.24 ± 0.45 | 3.92 ± 0.3 | 3.21 ± 0.06 |
| CL 66:2 | 20.44 ± 1.19 | **23.41 ± 0.93** | 19.55 ± 0.27 |
| CL 67:2 | 7.73 ± 0.24 | 8.23 ± 0.18 | 8.35 ± 0.15 |
| CL 68:2 | 28.58 ± 0.38 | **33.01 ± 1.26** | 27.85 ± 1.05 |
| CL 69:2 | 11.68 ± 0.34 | 11.84 ± 0.36 | **13.43 ± 0.45** |
| CL 70:3 | 12.73 ± 1.66 | **7.8 ± 0.68** | 12.40 ± 0.82 |
| CL 71:3 | 3.92 ± 0.52 | **1.49 ± 0.18** | 4.33 ± 0.47 |
| CL 72:4 | 1.67 ± 0.61 | 0.59 ± 0.1 | 1.75 ± 0.44 |
| Diacylglycerol | | | |
| DG 32:0 | 32.19 ± 1.88 | **44.95 ± 0.76** | 34.96 ± 5.08 |
| DG 34:1 | 49.53 ± 1.36 | **43.31 ± 0.96** | 47.95 ± 4.87 |
| DG 34:2 | 18.28 ± 1.26 | **11.74 ± 0.5** | 17.1 ± 0.95 |
| Diglucodiacylglycerol | | | |
| DGDG 30:0 | 2.89 ± 0.12 | **4.67 ± 0.18** | 3.24 ± 0.2 |
| DGDG 30:1 | 2.28 ± 0.06 | **1.15 ± 0.08** | 1.99 ± 0.1 |
| DGDG 32:0 | 11.95 ± 0.71 | **20.64 ± 0.73** | **13.05 ± 1.31** |
| DGDG 34:1 | 67.32 ± 0.55 | **60.05 ± 0.77** | 66.84 ± 0.1 |
| DGDG 35:1 | 6.93 ± 0.18 | **10.35 ± 0.34** | 6.9 ± 0.31 |
| DGDG 36:2 | 8.63 ± 0.36 | **3.14 ± 0.095** | 8.0 ± 0.51 |

| Stationary Phase | OG117 | OG117Δ*dak* | OG117Δ*dak*::*dak* |
|---|---|---|---|
| Phosphatidylglycerol | | | |
| PG 30:0 | 11.72 ± 0.28 | **7.24 ± 0.24** | 11.09 ± 0.16 |
| PG 32:0 | 11.09 ± 0.46 | 10.42 ± 0.55 | 11.11 ± 0.35 |
| PG 34:1 | 30.91 ± 0.9 | **37.78 ± 0.87** | 31.59 ± 0.56 |
| PG 35:1 | 38.47 ± 0.48 | **39.68 ± 1.14** | 38.45 ± 0.56 |
| PG 36:1 | 2.99 ± 0.07 | **1.94 ± 0.05** | 3.07 ± 0.06 |
| PG 37:2 | 4.72 ± 0.13 | **2.9 ± 0.15** | 4.6 ± 0.12 |
| PG 38:2 | 0.09 ± 0.005 | 0.05 ± 0.001 | 0.09 ± 0.008 |
| Lysylphosphatidylglycerol | | | |
| LysylPG 30:0 | 13.13 ± 0.38 | **5.54 ± 0.18** | 12.43 ± 0.37 |
| LysylPG 30:1 | 5.95 ± 0.24 | **3.91 ± 0.29** | 5.33 ± 0.15 |
| LysylPG 32:0 | 8.7 ± 0.49 | **5.27 ± 0.37** | 9.09 ± 0.38 |
| LysylPG 32:1 | 14.25 ± 0.5 | **16.5 ± 0.58** | 13.77 ± 0.31 |
| LysylPG 34:1 | 24.79 ± 0.71 | **33.82 ± 0.69** | **26 ± 0.25** |
| LysylPG 34:2 | 1.55 ± 0.19 | **3.24 ± 0.36** | 1.66 ± 0.26 |
| LysylPG 35:1 | 29.91 ± 1.56 | 29.31 ± 0.6 | 29.84 ± 0.37 |
| LysylPG 36:2 | 1.75 ± 0.18 | 2.42 ± 0.3 | 1.88 ± 0.31 |
| Cardiolipin | | | |
| CL 64:2 | 13.96 ± 0.44 | **10.51 ± 0.36** | **13.05 ± 0.91** |
| CL 65:2 | 11.32 ± 0.27 | **8.91 ± 0.29** | **10.08 ± 0.86** |
| CL 66:2 | 17.25 ± 0.4 | 16.66 ± 0.36 | 17.27 ± 0.23 |
| CL 67:2 | 13.37 ± 0.3 | **14.42 ± 0.27** | 12.89 ± 0.45 |
| CL 68:2 | 17.82 ± 0.63 | **19.27 ± 0.34** | **18.94 ± 1.15** |
| CL 69:2 | 12.58 ± 0.29 | **14.14 ± 0.28** | 13.05 ± 0.6 |
| CL 70:3 | 9.96 ± 0.25 | **12.25 ± 0.3** | 10.56 ± 0.28 |
| CL 71:3 | 2.9 ± 0.2 | 2.95 ± 0.15 | 3.27 ± 0.17 |
| CL 72:4 | 0.84 ± 0.06 | 0.89 ± 0.18 | 0.9 ± 0.1 |
| Diacylglycerol | | | |
| DG 32:0 | 41.14 ± 0.3 | **43.88 ± 1.38** | **45.12 ± 2.08** |
| DG 34:1 | 43.64 ± 0.6 | 43.35 ± 1.38 | **39.16 ± 1.19** |
| DG 34:2 | 15.22 ± 0.82 | **12.78 ± 0.25** | 15.72 ± 0.9 |
| Diglucodiacylglycerol | | | |
| DGDG 30:0 | 5.25 ± 0.23 | 5.91 ± 0.1 | 5.44 ± 0.24 |
| DGDG 30:1 | 2.47 ± 0.03 | 1.61 ± 0.2 | 2.38 ± 0.04 |
| DGDG 32:0 | 12.04 ± 0.31 | **13.17 ± 0.18** | 12.44 ± 0.62 |
| DGDG 34:1 | 59.03 ± 0.9 | **51.66 ± 0.57** | 58.95 ± 1.23 |
| DGDG 35:1 | 13.85 ± 0.88 | **23.08 ± 0.64** | 13.7 ± 0.78 |
| DGDG 36:2 | 7.36 ± 0.24 | **4.57 ± 0.14** | 7.1 ± 0.38 |

**Supplemental Table 2. Membrane lipid composition of E. faecalis OG117 strains.** Percent fatty acyl chain composition for each lipid species. Bold values indicate significant difference as compared to OG117 ($p \leq 0.05$, two-way ANOVA, with Tukey's test for multiple comparisons).

**Supplemental Table 3. Strains and plasmids used in this study.**

| Strain/Plasmid | Relevant Characteristics | Ref |
|---|---|---|
| *E. faecalis* | | |
| OG1RF | Laboratory strain of *E. faecalis*, WGS accession CP002621.1 | [1] |
| OG1RFΔ*madR* | Non-polar deletion of *madR* in OG1RF wild type background | [2] |
| OG1RF*madSA202E* | OG1RF with exchange of the *madS* gene for the allele with a C>A nucleotide change at position 605, resulting in the amino acid change A202E | This study |
| OG1RFΔ*liaR* | Non-polar deletion of *liaR* in OG1RF wild type background | [3] |
| OG1RFΔ*liaRmadSA202E* | OG1RFΔ*liaR* with exchange of the *madS* gene for the allele with a C>A nucleotide change at position 605, resulting in the amino acid change A202E | This study |
| OG117 | Derivative of OG1RF with the *E. faecalis* ATCC 4200 CRISPR1 *cas9* gene inserted in a neutral genomic insertion site | [4] |
| OG117Δ*dak* | Derivative of OG117 with a deletion of dak gene (OG1RF_12374) leaving only first 13 amino acids and 6 amino acids just prior to stop codon (Met N V T E I S A G Q F Q E V F V M K K Stop) | This study |
| OG117Δ*dak::dak* | Complementation of OG117Δ*dak* with the full length dak gene in the native chromosomal location | This study |
| OG117Δ*dakmadSA202E* | Derivative of OG117Δ*dak* with *madSA202E* allele, contains ser->tyr change in FabT at amino acid 36 in the DNA binding domain | This study |
| OG117Δ*dakmadSA202E*Δ*madLM* | Derivative of OG117Δ*dakmadSA202E* with deletion of *madLM* (OG1RF_11656 and OG1RF_11657) encoding the MadLM ABC transporter, contains ser->tyr change in FabT at amino acid 36 in the DNA binding domain | This study |
| OG117Δ*dakmadSA202E*Δ*madEFG* | Derivative of OG117Δ*dakmadSA202E* with deletion of *madEFG* (OG1RF_12267, OG1RF_12268, and OG1RF_12269), contains ser->tyr change in FabT at amino acid 36 in the DNA binding domain | This study |
| OG117Δ*dakmadSA202E*Δ*madEFG*::*madEFG* | Derivative of OG117Δ*dakmadSA202E*Δ*madEFG* with complementation of *madEFG* in the native chromosomal location, contains ser->tyr change in FabT at amino acid 36 in the DNA binding domain | This study |
| OG117Δ*dakmadSA202E*Δ*madEFG*Δ*madLM* | Derivative of OG117Δ*dakmadSA202EmadEFG* with deletion of *madLM*, contains ser->tyr change in FabT at amino acid 36 in the DNA binding domain | This study |
| OG117Δ*dakmadSA202E*Δ*dltA* | Derivative of OG117Δ*dakmadSA202E* with deletion of *dltA* (OG1RF_12112), contains | This study |

| | | |
|---|---|---|
| | ser->tyr change in FabT at amino acid 36 in the DNA binding domain | |
| OG117Δ*dakmadSA202E*Δ*dltA*::*dltA* | Derivative of OG117Δ*dakmadSA202EdltA* with complementation of *dltA* in the native chromosomal location, contains ser->tyr change in FabT at amino acid 36 in the DNA binding domain | This study |
| OG117Δ*liaX* | Derivative of OG117 with deletion of the *liaX* gene | [5] |
| OG117Δ*liaX*Δ*madEFG* | Derivative of OG117Δ*liaX* with deletion of *madEFG* | This study |
| OG117Δ*liaX*Δ*madEFG::madEFG* | Derivative of OG117Δ*liaX*Δ*madEFG* with complementation of *madEFG* in the native chromosomal location | This study |
| OG117Δ*liaX*Δ*dltA* | Derivative of OG117Δ*liaX* with deletion of *dltA* | This study |
| OG117Δ*liaX*Δ*dltA::dltA* | Derivative of OG117Δ*liaX*Δ*dltA* with complementation of *dltA* in the native chromosomal location | This study |
| Plasmids | | |
| pHOU1 | Derivative of pCJK47 in which the *erm*(C) gene was replaced by *aph-2"-ID*; confers GEN resistance | [6] |
| pCE | *oriT* from pCF10, containing the constitutive P$_{bacA}$ promoter from pPD1 to express the guide RNA, *cat* for chloramphenicol selection, and *pheS** for p-chloro-phenylalanine counterselection | [4] |
| pAT392 | *oriR*$_{\mathrm{pAM}\beta1}$, *oriR*$_{\mathrm{pUC}}$ *oriT*$_{\mathrm{RK2}}$ *spc lacZ*α P2 *aac(6')-aph(2")* | [7] |
| pAT392::*madR* | Derivative of pAT392 with *madR* and upstream sequence containing ribosomal binding site | [2] |

**Supplemental Table 4. Primers used in this study.**

| Primer | Sequence | Notes |
|---|---|---|
| dltA_spacer_F | GTAATTAATATGATTCAAACGATTGATGAAgtt ttagagtcatgttgtttagaatgg | forward primer for *dltA* targeting guide RNA spacer |
| dltA_spacer_R | TTCATCAATCGTTTGAATCATATTAATTACtttc attgctattatacccatgtag | reverse primer for *dltA* targeting guide RNA spacer |
| pCE_dltA_AF | atattacagctccagatccatatccttcttCCGAAACTTG GCGGCTAAAT | Forward primer for cloning upstream crossover region of *dltA* into pCE |
| dltA_linker_AR | GCCGCAATTAGCAGAACGTATAGCCGCCTCCT TAAAACTC | Reverse primer for cloning upstream crossover region of *dltA* into pCE |
| dltA_linker_BF | GAGGCGGCTATACGTTCTGCTAATTGCGGCCT TG | Forward primer for cloning downstream crossover region of *dltA* into pCE |
| pCE_dltA_BR | gaagcgaaaaaggagaagtcggttcagaaaCCAACTT AGGAACGGTTTGTTAAAG | Reverse primer for cloning downstream crossover region of *dltA* into pCE |
| dltA_Ext_F | CAAATGCCTAACGAAGAGGC | Screening primer for *dltA* mutants, anneals to gDNA external to crossover region upstream of the gene |
| dltA_Int_F | CTGTAAAAGCCGTTTTGAAGC | Screening primer for *dltA* mutants, anneals to gDNA internal to crossover region upstream of the gene |
| dltA_Int_R | CCCTAAAAAGCCAAGTAAGGTTG | Screening primer for *dltA* mutants, anneals to gDNA internal to crossover region downstream of the gene |
| dltA_Ext_R | GATAAGGTCATGTGCCAACG | Screening primer for *dltA* mutants, anneals to gDNA external to crossover region downstream of the gene |
| yxdL_spacer_F | TTTATTGGCAACAATCGATAGTCCAACAGAgtt ttagagtcatgttgtttagaatgg | forward primer for *madL* targeting guide RNA spacer |
| yxdL_spacer_R | TCTGTTGGACTATCGATTGTTGCCAATAAAtttc attgctattatacccatgtag | reverse primer for *madL* targeting guide RNA spacer |
| pCE_yxdLM_AF | atattacagctccagatccatatccttcttAGAAGAAAG AGCGTGCTTTA | Forward primer for cloning upstream crossover region of *madLM (yxdLM)* into pCE |
| yxdLM_linker_AR | GCTTAACGATTTTTTATAAATTTTCCACTCCTA TTCTTCTC | Reverse primer for cloning upstream crossover region of *madLM (yxdLM)* into pCE |

| | | |
|---|---|---|
| yxdLM_linker_BF | GGAGTGGAAAATTTATAAAAAATCGTTAAGCATGCAC | Forward primer for cloning downstream crossover region of *madLM (yxdLM)* into pCE |
| pCE_yxdLM_BR | gaagcgaaaaaggagaagtcggttcagaaaCGAACTTCGTTGTTAGGAGC | Reverse primer for cloning downstream crossover region of *madLM (yxdLM)* into pCE |
| yxdLM_EF | CTTAGTAGATGAATTGGAAGCAC | Screening primer for *madLM (yxdLM)* mutants, anneals to gDNA external to crossover region upstream of the gene |
| yxdLM_IF | CACCTTTCACGGTTATATCTAATCG | Screening primer for *madLM (yxdLM)* mutants, anneals to gDNA internal to crossover region upstream of the gene |
| yxdLM_IR | CAGATAATCCATATTCTTGACGC | Screening primer for *madLM (yxdLM)* mutants, anneals to gDNA internal to crossover region downstream of the gene |
| yxdLM_ER | CCTGCACAAATGGTTAAGGC | Screening primer for *madLM (yxdLM)* mutants, anneals to gDNA external to crossover region downstream of the gene |
| EF2987_spacer_F | TCAGGCCAGAAGAAGCGAAATAAAAAAGTGgttttagagtcatgttgtttagaatgg | forward primer for *madG* targeting guide RNA spacer |
| EF2987_spacer_R | CACTTTTTTATTTCGCTTCTTCTGGCCTGAtttcattgctattatacccatgtag | reverse primer for *madG* targeting guide RNA spacer |
| pCE_EF2987_AF | atattacagctccagatccatatccttcttTGCAGCGATGAACCTTGTAGA | Forward primer for cloning upstream crossover region of *madEFG* into pCE |
| EF2987_linker_AR | GTTTCTTTGCTGTTTATATTTATTTGTTCCTCCTTGATTTCTGTTGAC | Reverse primer for cloning upstream crossover region of *madEFG* into pCE |
| EF2987_linker_BF | GAGGAACAAATAAATATAAACAGCAAAGAAACAGCCATTTTTG | Forward primer for cloning downstream crossover region of *madEFG* into pCE |
| pCE_EF2987_BR | gaagcgaaaaaggagaagtcggttcagaaaTTCTGCGGCTGTTAGCTTTATTC | Reverse primer for cloning downstream crossover region of *madEFG* into pCE |
| EF2987_Ext_F | GAGTTTACCTATGCGCCGCC | Screening primer for *madEFG* (EF2987-EF2985) mutants, anneals to gDNA external to crossover region upstream of the gene |
| EF2987_Int_F | GTGCGTTTCTCAGTCAACAAGG | Screening primer for *madEFG* (EF2987-EF2985) mutants, anneals to gDNA internal to crossover region upstream of the gene |

| | | |
|---|---|---|
| EF2987_Int_R | CAGTCATTCCTGTAATCCCCGA | Screening primer for *madEFG* (EF2987-EF2985) mutants, anneals to gDNA internal to crossover region downstream of the gene |
| EF2987_Ext_R | GAGGTCATCCTGTGATGGTG | Screening primer for *madLM madEFG* (EF2987-EF2985) mutants, anneals to gDNA external to crossover region downstream of the gene |
| DAK_comp_spacer_F | CGTTTTTATTTTTTCATCACGAAAACTTCCgttttagagtcatgttgtttagaatgg | forward primer for *dak* targeting guide RNA spacer |
| DAK_comp_spacer_R | GGAAGTTTTCGTGATGAAAAAATAAAAACGtttcattgctattatacccatgtag | reverse primer for *dak* targeting guide RNA spacer |
| delDAK2_DownF_NotI | GGGCGGCCGCAAGCGATCCACGTCATATTG | Forward primer for cloning downstream crossover region of *dak* into pCE, NotI restriction site underlined |
| delDAK2_DownR | GTGTATCCATACTTATTCTCAGCAGAATAG | Reverse primer for cloning downstream crossover region of *dak* into pCE |
| delDAK2_UpF | TTCCTGGAACTGACCTGC | Forward primer for cloning upstream crossover region of *dak* into pCE |
| delDAK2_UpR_PstI | GGCTGCAGCATTCGGTCTGTAGATATGGC | Reverse primer for cloning upstream crossover region of *dak* into pCE, PstI restriction site underlined |
| delDAK2_ExtDownF | GCATATGCTTCGGACTTCGC | Screening primer for *dak* mutants, anneals to gDNA external to crossover region downstream of the gene |
| delDAK2_IntDownF | TCGAAACGTTAACCGGCTCT | Screening primer for *dak* mutants, anneals to gDNA internal to crossover region downstream of the gene |
| delDAK2_IntUpR | TCGCAGTGGATGTCTACACG | Screening primer for *dak* mutants, anneals to gDNA internal to crossover region upstream of the gene |
| delDAK2_ExtUpR | TAAAGTGGAAGCGCTGGACTA | Screening primer for *dak* mutants, anneals to gDNA external to crossover region upstream of the gene |
| BamHI_2987Comp_up_F | GACGGATCCGATCCTGTGAACATGTTAGGC | Forward primer for cloning *madEFG* complementation into pHOU1, BamHI restriction site underlined |

| | | |
|---|---|---|
| EcoRI_2987Comp_Down_R | GACGAATTCCTATACAAGACAGCCAGTAGTGC | Reverse primer for cloning *madEFG* complementation into pHOU1, EcoRI restriction site underlined |
| BamHI_2049Comp_Up_F | GACGGATCCAGAAGAAAGAGCGTGCTTTA | Forward primer for cloning *madLM* complementation into pHOU1, BamHI restriction site underlined |
| BamHI_2049Comp_Down_R | CAGGGATCCCGAACTTCGTTGTTAGGAGC | Reverse primer for cloning *madLM* complementation into pHOU1, BamHI restriction site underlined |
| dltA_EcoRI_F | GACGAATTCCCGAAACTTGGCGGCTAAAT | Forward primer for cloning *dltA* complementation into pHOU1, EcoRI restriction site underlined |
| dltA_BamHI_R | GACGGATCCCCAACTTAGGAACGGTTTGTTAAAG | Reverse primer for cloning *dltA* complementation into pHOU1, BamHI restriction site underlined |
| 2050_F | GGTGACCCATGATCCGCTAG | qRT-PCR primer, *madL* |
| 2050_R | TCGACACCTTCAATCTCAGCC | qRT-PCR primer, *madL* |
| 2752_F | ACTGGCTCGCTGGATTCAAA | qRT-PCR primer, *madA* |
| 2752_R | TGTACGGGTCATGTGTCACC | qRT-PCR primer, *madA* |
| 2749_F | GCGCATCGAAAGCTGGTTTT | qRT-PCR primer, *dltA* |
| 2749_R | ACTTCGGTGGCTAACTCAGG | qRT-PCR primer, *dltA* |
| 2987_F | CATCGCTGTACCGCAAAAGC | qRT-PCR primer, *madG* |
| 2987_R | TTTTCGCTTTGCCCGCTTTG | qRT-PCR primer, *madG* |
| gyrB_F | AAAAGGCATGTTGGCTTCAAA | qRT-PCR primer, DNA gyrase housekeeping gene |
| gyrB_R | GCTTCCCTGGCAAGTTGCTA | qRT-PCR primer, DNA gyrase housekeeping gene |
